# De novo antibody discovery in human blood from full-length single B cell transcriptomics and matching haplotyped-resolved germline assemblies

**DOI:** 10.1101/2024.03.26.586834

**Authors:** John Beaulaurier, Lynn Ly, J. Andrew Duty, Carly Tyer, Christian Stevens, Chuan-tien Hung, Akash Sookdeo, Alex W. Drong, Shreyas Kowdle, Daniel J. Turner, Sissel Juul, Scott Hickey, Benhur Lee

**Affiliations:** Oxford Nanopore Technologies, Inc. New York, New York 10013, USA; Icahn School of Medicine at Mount Sinai, New York, New York 10029, USA

## Abstract

Immunoglobulin (IGH, IGK, IGL) loci in the human genome are highly polymorphic regions that encode the building blocks of the light and heavy chain IG proteins that dimerize to form antibodies. The processes of V(D)J recombination and somatic hypermutation in B cells are responsible for creating an enormous reservoir of highly specific antibodies capable of binding a vast array of possible antigens. However, the antibody repertoire is fundamentally limited by the set of variable (V), diversity (D), and joining (J) alleles present in the germline IG loci. To better understand how the germline IG haplotypes contribute to the expressed antibody repertoire, we combined genome sequencing of the germline IG loci with single-cell transcriptome sequencing of B cells from the same donor. Sequencing and assembly of the germline IG loci captured the IGH locus in a single fully-phased contig where the maternal and paternal contributions to the germline V, D, and J repertoire can be fully resolved. The B cells were collected following a measles, mumps, and rubella (MMR) vaccination, resulting in a population of cells that were activated in response to this specific immune challenge. Single-cell, full-length transcriptome sequencing of these B cells resulted in whole transcriptome characterization of each cell, as well as highly-accurate consensus sequences for the somatically rearranged and hypermutated light and heavy chain IG transcripts. A subset of antibodies synthesized based on their consensus heavy and light chain transcript sequences demonstrated binding to measles antigens and neutralization of measles live virus.

## Introduction

The immune system is a complex network of cells, tissues and organs that work together to defend the body against infections and foreign substances. It is broadly categorized into two main components: the innate immune system, which responds quickly to infections but does not exhibit specificity or memory, and the adaptive immune system, which provides a specific, targeted response to pathogens, and which develops a memory of past exposures. The adaptive immune system is composed of T cells, which create a cell-mediated response to pathogen exposure, and B cells, which produce antibodies. To recognize a wide variety of pathogens, both T and B cells must be able to produce highly diverse antigen receptors. This is achieved by extensive recombination of their receptor-gene segments, creating a unique immune repertoire in every individual (Tonegawa 1983; Watson et al. 2017). In B cell maturation, multiple distinct subtypes of cells arise during an immune response. These include plasma cells and plasmablasts, which are transient antibody-secreting populations appearing during the acute phase of an immune response, and memory B cells, which produce low levels of antibodies in order to ‘remember’ a specific antigen and can quickly differentiate into plasmablasts upon re-exposure to a previously encountered antigen. As a result, re-exposure to the same or closely related pathogen or antigen results in clonal expansion from the circulating memory B cell population, enabling a more rapid and robust immune response.

Sequencing immune-gene transcripts of individuals within a population reveals allelic diversity and allows comparison between immune responses to pathogenic antagonists, cancer, and autoimmune diseases (Hansen et al. 2024; Rodriguez et al. 2023; Boyd et al. 2010). Single-cell transcriptome sequencing of immune cells enables closer examination of heterogeneity within these cell populations and has led to a greater understanding of immune cell gene expression, heavy-light chain pairing, VDJ recombination, somatic hypermutation, and class-switch recombination than can be derived from bulk analyses (Papalexi and Satija 2018; Di Noia and Neuberger 2007; Dudley et al. 2005; Jaffe et al. 2022). Single-cell sequencing using long-read approaches can produce reads that span entire transcripts, removing the need for isoform reconstruction and is therefore preferable to short read approaches that can miss novel transcripts and isoforms (Singh et al. 2019; Subas Satish et al. 2022; Engblom et al. 2023). This is particularly important in the context of antibody sequencing because the variable domain must be unambiguously paired with the constant domain in order to determine both the VDJ recombination and the class-switch recombination in the same antibody transcript. Long-read, single-cell transcriptomes of B cells paired with high-quality germline genome assembly enables unambiguous annotation of allele specific transcripts after recombination. Additionally, germline immunoglobulin assembly makes it possible to filter antibody clones based on the presence or absence of specific alleles, for example variable alleles, that are present within an individual versus ruling out closely matched alleles that are not present within the individual’s germline sequence.

While single-cell methods have provided novel insights into the transcriptional and post-transcriptional processes involved in the immune response, the genetic components have been harder to unravel. This is due to both the high levels of polymorphism and the structural complexity of the loci involved (Nurk et al. 2022; Rodriguez et al. 2020; Watson et al. 2013; Rodriguez et al. 2023). The B cell receptor (BCR) is composed of immunoglobulin (IG) heavy and light chains. The heavy chain is encoded in humans by the IGH locus on chromosome 14 and the light chain is encoded by either the IG kappa (IGK) locus on chromosome 2 or the IG lambda (IGL) locus on chromosome 22. The IG loci have complex genomic organization, high sequence diversity and undergo extensive rearrangement in B cells during antigen-mediated activation and maturation. Consequently, assembly of these loci benefits from use of long reads that can resolve their complex structures and improve haplotype phasing (Kolmogorov et al. 2023). However, individual single-cell transcriptomes have so far not been paired with germline assemblies from the same donor. Haplotyped structural variants and heterogeneity of immunoglobulin V, D, J, and C gene segments may affect the antibody repertoire of a given individual, representing the potential of future precision medicine based on individual variation (Kidd et al. 2016; Kenter et al. 2021; Peres et al. 2023).

In the work presented here, we generated full-length single-cell antibody transcripts from memory B cells (MBCs) and antibody-secreting cells (ASCs), as well as haplotype-resolved germline assemblies of IG loci from a single donor (Fig. 1). The assemblies were annotated against IMGT/GENE-DB (Giudicelli et al. 2022), a public database of immunoglobulin V, D, J, and C sequences, and the transcripts were mapped against this personalized set of IG genes. Our approach allows some allelic annotations to be filtered out if they are not present in the matched germline locus, as those reference transcripts could not have been translated into an antibody in that individual. Finally, we validated functional antibody clones against viral antigens and characterized the clonal diversity at the single-cell level of paired heavy and light chain clones.

**Figure 1.**
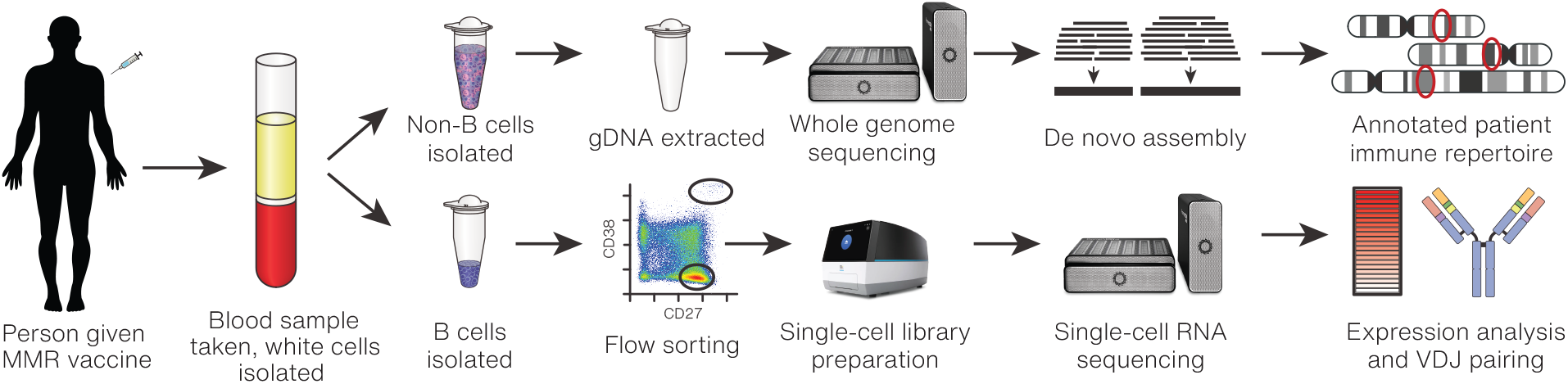
Experimental design schematic. Peripheral blood mononuclear cells (PBMCs) collected from a donor at 6 days post MMR vaccination were separated into B cells for single-cell sequencing and monocytes for germline sequencing and assembly.

## Results

### A donor-specific IGH assembly is highly contiguous and fully phased

Understanding the germline IG allelic composition is helpful for precise annotation and validation of expressed clonal antibody sequences. However, haplotyped IG assemblies have remained difficult to generate due to the high sequence homology of variants and complex genomic structure of each locus, notably the segmental duplications in IGK (Lefranc 2001b). To assess the full repertoire of donor-specific germline immune alleles, we depleted B and T cells from whole blood and extracted DNA for whole genome nanopore sequencing (Methods). Reads were *de novo* assembled and phased, resulting in a highly contiguous personalized whole genome assembly (statistics in Supplemental Table S1). The IGH, IGL, and IGK loci were isolated from this assembly for in-depth annotation and study.

The 2 Mb IGH region was contained within a 79 Mb contig and separated into two phase blocks. The phase block gap did not contain heterozygous SNPs but was manually resolvable using a heterozygous 172 bp deletion between IGHV3-20 and the pseudogene IGHV3-19, resulting in a fully phased IGH region (Fig. 2A). Additionally, a 37 kb alternate contig representing the known 5-10-1 / 3-64D structural variant (Lefranc 2001a) was manually placed into a haplotype. All IGHJ and constant gene segments were present in the expected order. A block of IGHD gene segments was missing from each haplotype; haplotype 1 was missing D3-3 through D2-8, and haplotype 2 was missing D6-6 through D5-1.

**Figure 2.**
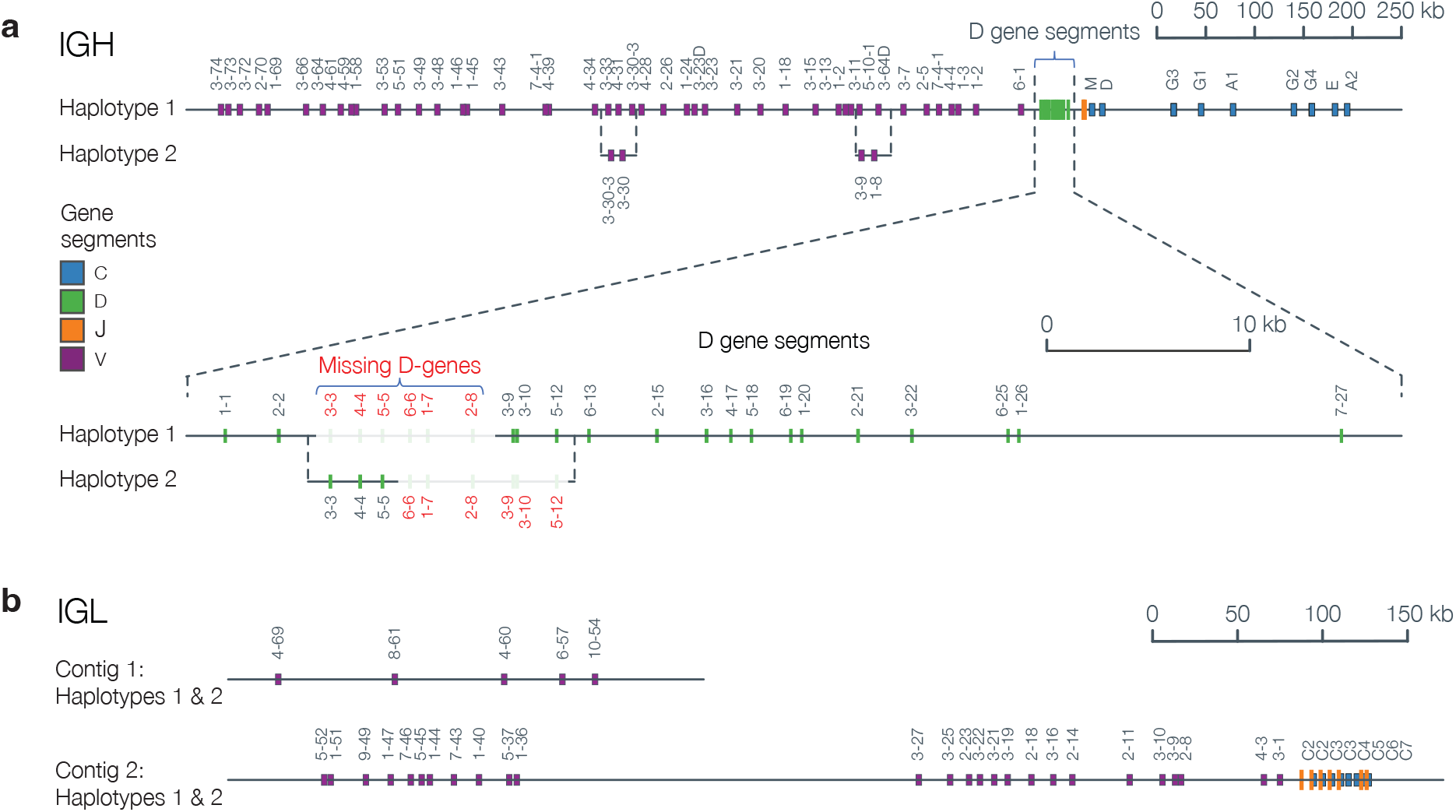
A donor-specific, haploid assembly was created using Flye and HapDup (Methods). Colored bars indicate the position on the contig of functional V, D, J, and C gene segments. (*A*) The fully phased IGH locus, with the dashed lines depicting alternative structures in sections where the assembly differs structurally between the two haplotypes, such as the IGHV3-9 & IGHV1-8 structural variant. The D gene cluster is expanded to show that a block of D gene segments is missing from each haplotype. (*B*) The IGL locus, where the IGL C V-CLUSTER was assembled into a separate contig from the remaining V-CLUSTERs and the J-C-CLUSTERs. All gene segments were shared between both haplotypes.

The IGL V clusters A and B, as well as the J-C cluster, were contained on a fully-phased 26 Mb contig; V cluster C was present on a separate phased contig (Fig. 2B). The IGL region typically contains 7 to 11 highly similar J-C cassettes, ranging from 3.5-5.6 kb, each containing a J and a C gene segment (Lefranc 2001c). This IGL J-C cluster contains 9 cassettes, including one novel IGLC3 allele that differed from the closest known alleles by a single SNV (Supplemental Fig. S1).

Although ∼660 kb of IGK sequence was assembled, it was relatively discontiguous (split into 7+ contigs). Furthermore, IGK could not be fully resolved due to two major challenges: 1) It consists of two highly similar ∼400 kb segmental duplications and 2) these segmental duplications are separated by 800 kb of highly repetitive sequence (Lefranc 2001b). Consequently, the assembler collapsed the homologous sequences between the two segmental duplications, and allele calling was not possible for collapsed genes.

### Single cell transcripts and isoforms are di<erentially expressed among B cell subpopulations and unambiguously annotated using nanopore sequencing

To understand the expression of immune genes in response to an immune challenge, our donor received a measles, mumps, and rubella (MMR) vaccination followed by a longitudinal series of blood draws. Daily peripheral blood mononuclear cell (PBMC) samples were monitored for the appearance of CD27++/CD38++ expressing ASCs by fluorescence-activated cell sorting (FACS; Supplemental Fig. S2). This putative population of ASCs appeared on day 6 post-vaccination, at which point we collected 80 mL of whole blood for B cell enrichment and sorting. We used a multi-parameter FACS strategy (Methods), involving doublet exclusion, viability gating, and a T/monocyte/NK cell exclusion step to ensure a highly pure population of CD19+/IgM-B cells for sorting into CD38++/CD27++ ASCs in the P7 gate (referred to henceforth as the ASC gate) and CD38-/CD27+ MBCs in the P8 gate (MBC gate). Approximately 1,000 and 30,000 cells were collected from the ASC and MBC gates, respectively, and were subsequently processed for single-cell transcriptome sequencing.

In order to characterize single-cell expression within the B cell subpopulations, the single-cell, whole transcriptome libraries of both the ASC gate and MBC gate were split in half and sequenced by both short-read and nanopore sequencing platforms. Single-cell cDNA sequences from the short-read and nanopore libraries of both the ASC and MBC gates were compared for cell barcode overlap and unique molecular identifier (UMI) counts per cell. UMI comparisons revealed a high degree of concordance between the two sequencing technologies in both the ASC gate (Supplemental Fig. S3) and MBC gate (Supplemental Fig. S4).

Because nanopore sequencing generates reads covering full-length transcripts, expression was tabulated not only at the gene-level, but also at the transcript-level in order to quantify isoform expression of each gene. When compared to the MBC gate cellular landscape provided by gene-level expression using short reads (Fig. 3A), the transcript-level expression provided by long read sequencing revealed a higher degree of cellular heterogeneity in the population (Fig. 3B). An analysis of differential transcript expression across cell clusters (Methods) revealed a total of 1785 transcripts whose expression in cells from one cluster was significantly higher or lower than in all cells in the remaining clusters (Fig. 3C; Supplemental Table S2).

**Figure 3.**
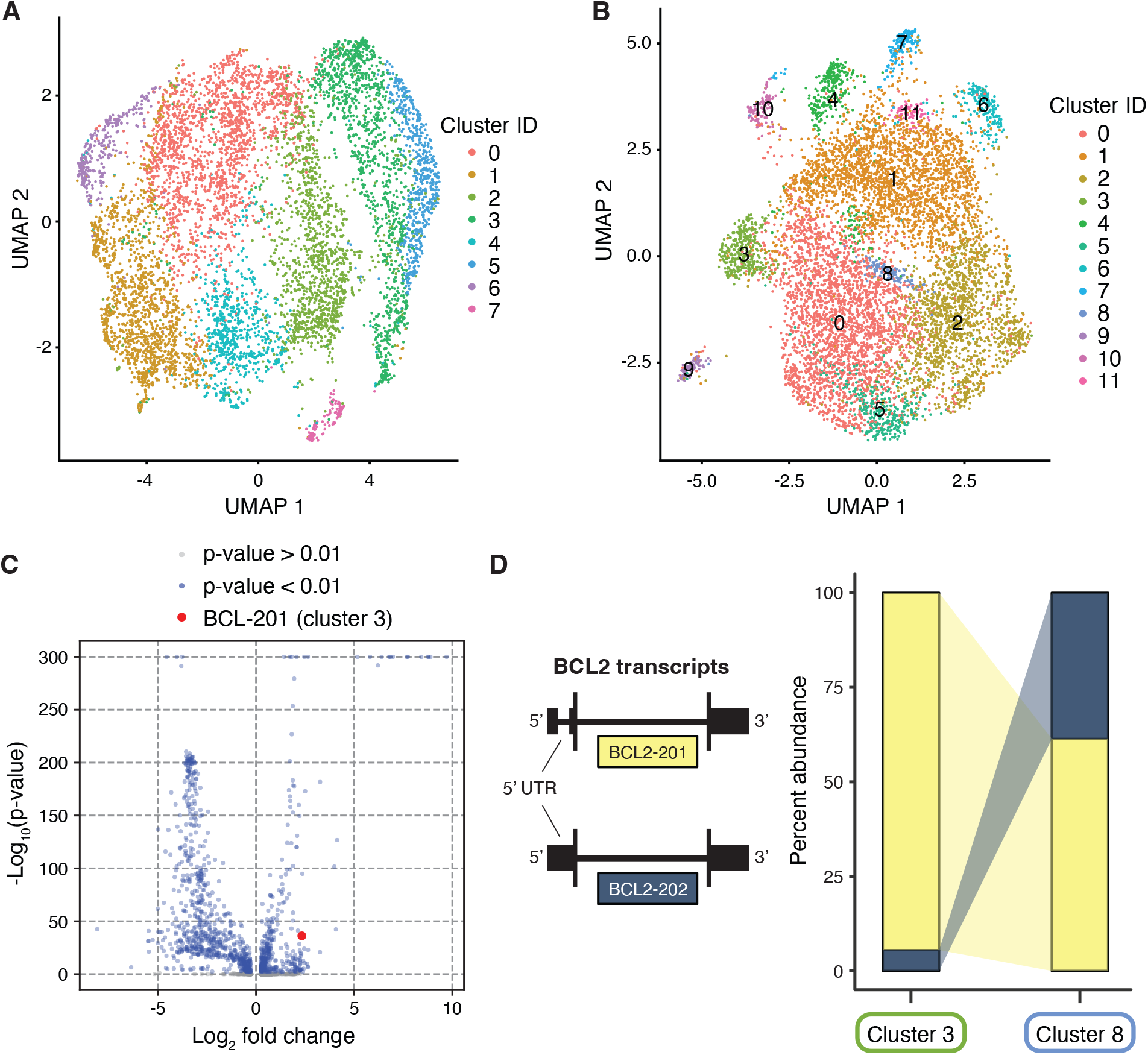
Single-cell expression data from the MBC gate. (*A*) UMAP visualization of cells clustered by gene expression quantified by short read sequencing. (*B*) UMAP visualization of cells clustered by isoform expression quantified by nanopore sequencing. (*C*) Volcano plot showing isoforms that are differentially expressed in one cluster relative to the entire population of cells. One differentially expressed isoform of the apoptosis regulating gene BCL2 is highlighted. (*D*) Relative abundance of two BCL2 transcripts in clusters 3 and 8 that differ in their 5’ UTR sequences. BCL2-202 isoforms make up only 5% of BCL2 transcripts in cluster 3 but represent 38% of BCL2 transcripts in cluster 8.

The differentially expressed transcripts can highlight how different isoforms of the same gene can be preferentially expressed in some cells, while other isoforms of the same gene are favored in other cells. For example, the gene BCL2 serves as a critical regulator of apoptosis and expresses nine known splice variants (Warren et al. 2019). The total expression of BCL2 in cluster 8 is comprised of similar levels of expression of two distinct isoforms: BCL2-201 (ENST00000333681.5) and BCL2-202 (ENST00000398117.1), which both express the same protein coding sequence but differ in the sequence of their 5’ UTRs. This is contrasted by cluster 3, where the vast majority of BCL2 gene expression derives from expression of the BCL2-202 isoform (Fig. 3D). The reason for this dramatic difference is unclear, but such striking examples of differential isoform expression across cell clusters are suggestive of important biological processes that would otherwise be obscured by gene-level analyses.

For the ASC gate, gene expression clustering of the cells based on nanopore sequencing revealed multiple cellular subpopulations, including CD38++ plasmablasts and plasma cells (Fig. 4A). Due to dynamic gating during the sorting process, some CD27+/CD38+ activated MBCs were also collected in the ASC gate, but these are easily identifiable as a separate population due to their distinct expression profiles (Fig. 4 A,B). Based on expression, cluster 2 appears to represent a population of non-proliferating plasma cells (MKI67-low, IRF4-high), while cells in cluster 4 are likely actively proliferating plasmablasts (MKI67-high, IRF4-low) (Fig 4B). Cells in both clusters 2 and 4 produce a high level of IGHC transcripts, consistent with their role in high production and secretion of antibodies. The remaining 374 CD38-cells in clusters 0, 1, and 3 are most likely residual MBCs that were missorted into the ASC gate, as they have with low expression of IGHC, DERL3, IRF4, and MKI67 but high expression of MS4A1/CD20 and HLA-DR. Nonetheless, our expression data were clearly able to differentiate these cells from plasmablasts or plasma cells.

**Figure 4.**
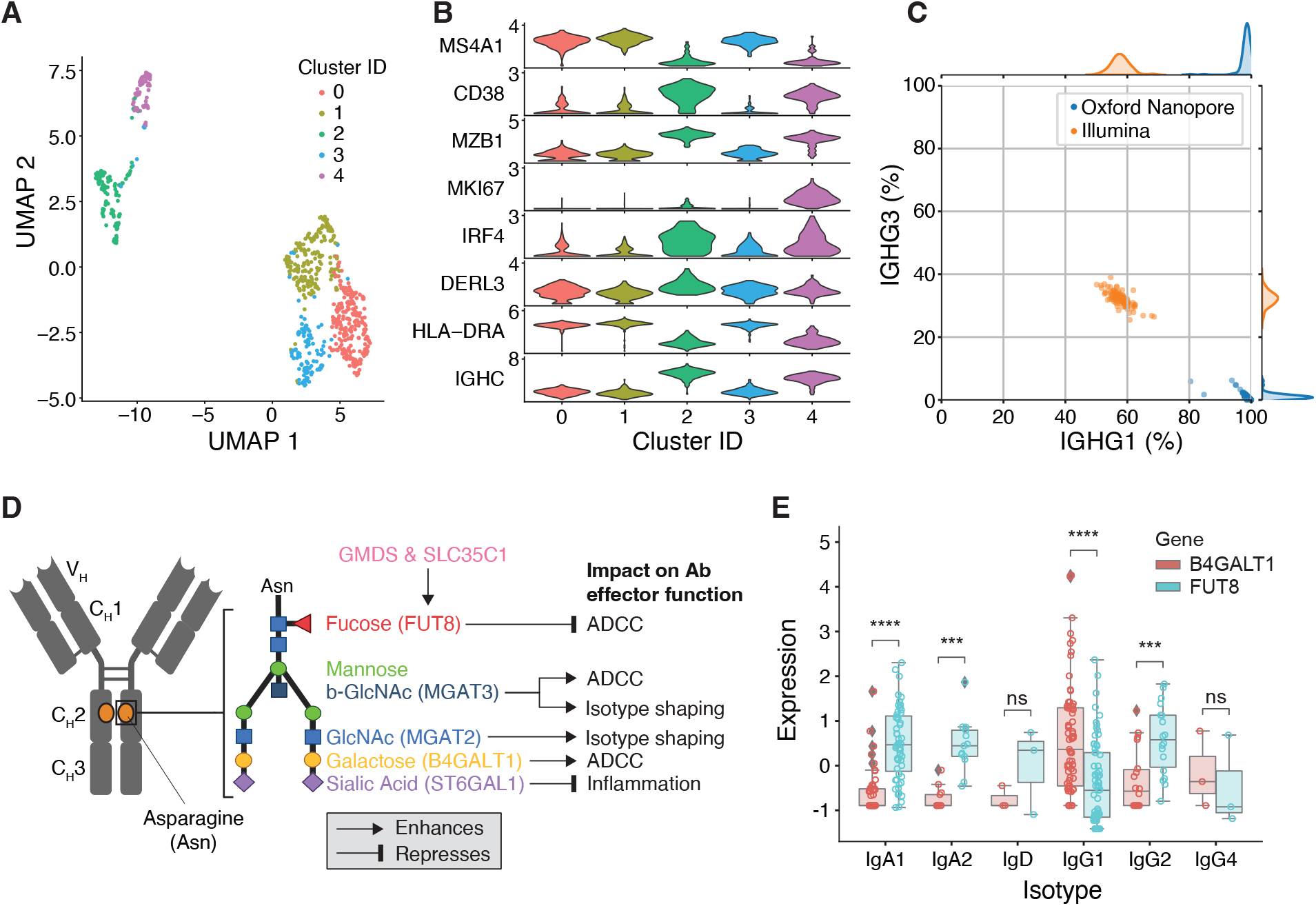
Single cell expression data from ASC gate. (*A*) UMAP visualization of cells clustered by gene expression quantified by nanopore sequencing. (*B*) Expression of marker genes differentiating in each cluster. (C) Proportion of reads assigned to IGHG1 and IGHG3 within each cell in both the short read and nanopore sequencing data sets. (*D*) Schematic illustrating the genes involved in glycosylation of the conserved asparagine residue in the CH2 domain of the heavy chain. (*E*) Differential expression of two genes in the glycosylation pathway, B4GALT1 and FUT8, in certain antibody isotypes of plasmablasts and plasma cells from the ASC gate.

Antibody specificity derives from the sequence of the complementarity determining regions (CDRs) of the light and heavy chains. The isotype of an antibody, such as IgA, IgD, IgE, IgG and IgM, is determined by the sequence of the constant domains in the heavy chain. IgG and IgA are further divided into subtypes that are determined largely by the sequence of the heavy chain hinge region and which activate the immune system in different ways. While each cell is expected to produce antibodies of a single isotype and subtype (e.g. IGHG1), the short-read sequencing data revealed ambiguous isotype assignments. Most ASCs had a mix of IGHG1 and IGHG3 assignments from short-reads, likely due to high sequence similarity in the limited portion of the constant domains that are captured in the fragmented short-read sequencing library. In contrast, the nanopore sequencing data showed no such ambiguous isotype and subtype assignments, with nearly all full-length transcript reads from each cell mapping definitively to either IGHG1 or IGHG3 (Fig. 4C).

### Antibody glycosylation phenotypes

Differential glycosylation of antibodies is known to affect their activities via its effects on Fc effector functions, serum half-life, and isotype shaping (Irvine and Alter 2020). GWAS studies have also linked SNPs in glycosyltransferases, such as ST6GAL1, B4GALT1, FUT8, and MGAT3, to IgG N-glycosylation phenotypes and certain isotypes (Shen et al. 2017; Wahl et al. 2018). Several core glycosylation genes are known to have specific impact on antibody-dependent cellular cytotoxicity (ADCC), isotype shaping, and inflammation (Fig. 4D). However, it is unclear if these genes are preferentially expressed at the level of single B cell clones.

We examined the expression of core antibody glycosylation genes known to affect antibody effector function in our expression data from both the ASC and MBC gates. FUT8 was significantly overexpressed relative to B4GALT1 in the majority of plasmablasts and plasma cells from the ASC gate that secreted IgA1 and IgA2 antibodies (Fig. 4E). The converse was true in plasmablast and plasma cells secreting IgG1 antibodies. Since increased fucosylation is associated with decreased ADCC function, our data suggest attenuation of ADCC activity in the secreted IgAs. The higher expression of B4GALT1 expression versus FUT8 expression in clones producing IgG1 suggest that these antibodies are likely to have higher ADCC activity.

While glycosylation gene expression patterns were not overtly associated with particular isotypes in the larger population of MBCs in the MBC gate, we found high MGAT3, MGAT2 and SLC35C1 expression to be restricted to a minority of MBCs (Supplemental Fig. S5). Furthermore, high MGAT3 and SLC35C1 expression in these MBC clones was mutually exclusive. This is consistent with the opposing roles of MGAT3 and SLC35C1 (via its enhancing effect on fucosylation) in increasing and decreasing ADCC activity, respectively.

### IG transcript enrichment and germline assembly mapping enables high-confidence consensus sequences for antibody synthesis

To first quantify the relative proportions of secreted and membrane-bound antibodies in each cell, we searched for reads containing the IGHC M exons, which reside on the 3’ end of the IGH transcript and harbor the transmembrane, connecting, and cytoplasmic tail regions. Although the majority of transcripts contained the secreted form of IGHC lacking the M exons, a subset of MBCs primarily expressed membrane-bound transcripts (Fig. 5A). In contrast, all of the ASCs primarily expressed the secreted forms.

**Figure 5.**
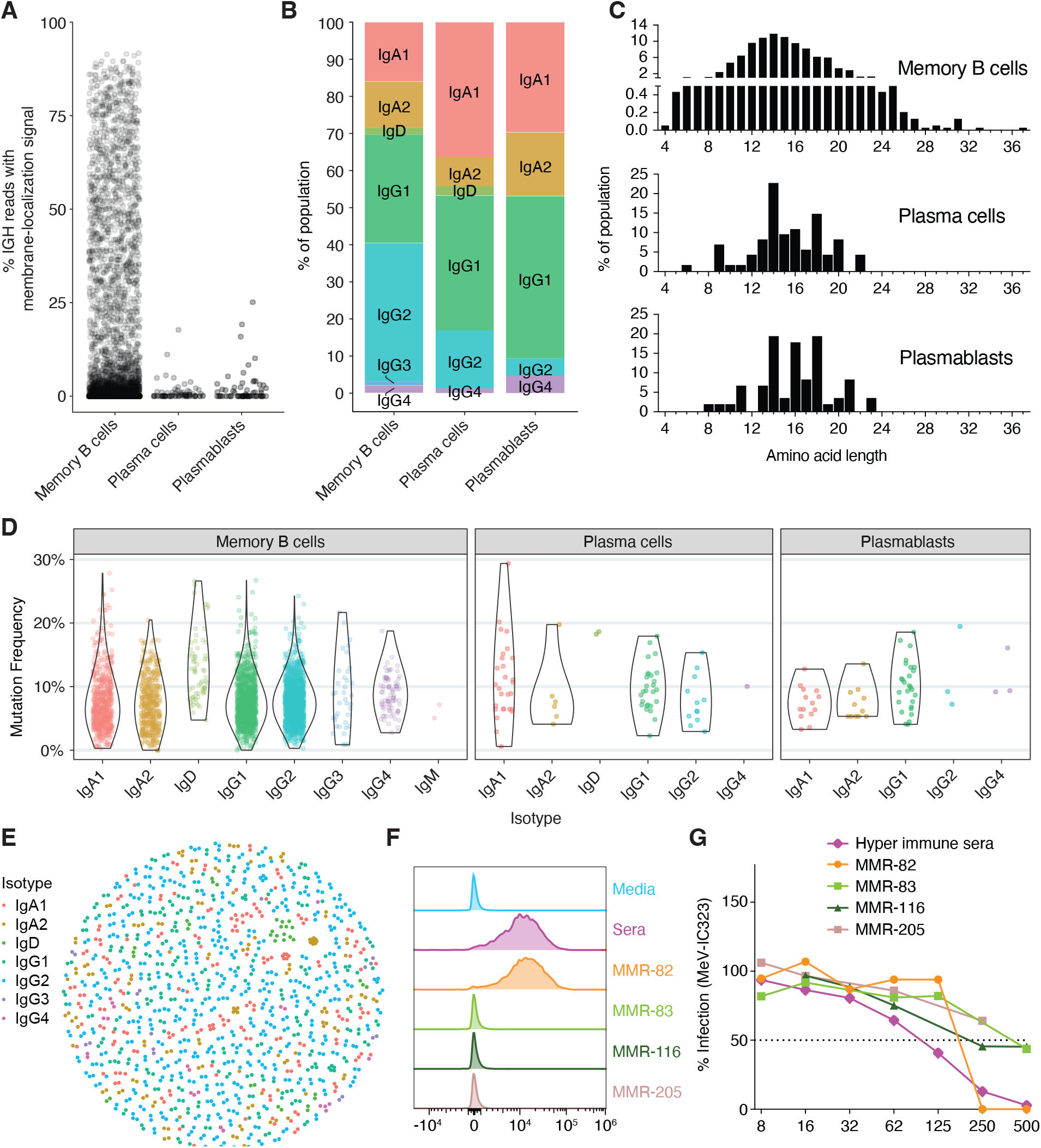
Characterizing single-cell immunoglobulin sequences and antibody synthesis. (*A*) The percentage of IGH reads in each cell possessing the M exons indicative of a membrane-bound antibody, stratified by cell type. (*B*) Proportions of each isotype identified from consensus IGH sequences, stratified by cell type. (*C*) CDR3 amino acid length distributions for all productive IGH consensus sequences, stratified by cell type. CDR3 lengths are enumerated between the junction anchor residues: Cys 104 and Try 118 (IMGT numbering) for each sequence. (*D*) Mutation frequency compared to the germline sequences in the FWR1-4 and CDR1-3 regions for each cell type and isotype. (*E*) Sequences clustered within each non-singleton clone by VDJ amino acid sequence similarity. (*F*) Putative positive recombinantly expressed antibodies sequenced from the ASC gate were screened by flow cytometry for binding to MeV surface expressed proteins (F or RBP/H) on virus infected Raji-DC-SIGN B cells (MOI 0.01). MMR-vaccinated, hyper-immune sera from the donor at the time of cell collection was used as a positive control. (*G*) Neutralization titers in Vero-hSLAM cells for each monoclonal antibody and hyper-immune sera in serial dilutions against a GFP expressing MeV-IC323 recombinant virus were calculated as percent infection compared to infection in cells alone.

Next, we paired heavy and light chain IG transcripts from each cell so that functional antibodies could be generated directly from the consensus sequences. The ASCs generated a large number of IG transcript reads from the whole transcriptome sequencing of each cell: on average 5319 from the light chain and 357 from the heavy chain.

The MBCs, however, had a much more diverse transcriptome that resulted in fewer IG transcript reads from each cell. To increase the IG transcript coverage for these cells, a second sequencing library was generated using biotin probe enrichment to enrich for IGH, IGL, and IGK-containing transcripts (Methods). This resulted in a 130-fold enrichment in the percentage of IG transcripts compared to the matching whole transcriptome sequencing dataset and made it possible to recreate high-accuracy heavy and light chain sequences from several thousand MBCs.

We created a custom pipeline (Supplemental Fig. S6) to produce high-accuracy heavy and light chain IG sequences for each cell (Methods). Briefly, IG transcript reads were identified and annotated for the V, J, and C gene segments using IgBLAST. Filtering based on V and C gene segment annotation quality metrics and positions in the read ensured that any low-quality or chimeric IG transcript reads were discarded. These remaining reads were grouped based on cell barcode and IG locus (IGH, IGK, or IGL), then medaka (https://github.com/nanoporetech/medaka) was used to generate a single consensus sequence per cell barcode and IG locus. The resulting heavy and light chain consensus sequences per cell were interrogated once more using IgBLAST to determine their isotype (Fig. 5B) and predict their ability to form productive and complete heavy or light chain proteins with in-frame junctions (CDR3s).

This pipeline generated 761 and 7653 consensus sequences that were predicted to be productive from the ASC and MBC gates, respectively. The length distributions varied by isotype and were consistent with expectations based on the literature and reference database (IMGT-GENE-DB) (Lefranc et al. 2005) (Supplemental Fig. S7).

Observed variation in the length of sequences from each isotype can be explained by the variable lengths of the CDR3 sequences for each subpopulation (Fig. 5C). A Gaussian-like distribution of CDR3 lengths in the memory population was consistent with average CDR3 lengths in peripheral MBC populations in healthy individuals, with a median of 14 amino acids (Duty et al. 2009; Tian et al. 2007; Dong et al. 2020). Interestingly we observed that 0.2% (8 sequences) of the MBC contained CDR3 lengths of 30aa or longer, including one CDR3 of 37aa, that were not selected in the ASC populations. Non-symmetrical distributions in the ASC populations, plasma cells, and plasmablasts show evidence of length biases, indicative of selection.

Nucleotide mutation frequencies observed in the V gene segment were within the standard range for post-germinal center, class-switched B cell populations with an average mutation frequency of 7.7% across all isotypes (Dong et al. 2020; Wrammert et al. 2008; Budeus et al. 2023) (Fig. 5D). Further, the mutation frequencies were determined by comparing the consensus transcript sequences to the personalized, annotated IG loci assemblies, where the V gene segment alleles have been confidently assigned. This likely resulted in a more accurate determination of mutation frequency than would be possible if relying on public databases of all V gene segment alleles.

Analysis of mutation by isotype revealed rarer IgD memory B cell and plasma cell populations (1% and 0.03% abundance within the two populations, respectively), which did not contain IgM transcripts, had high mutation frequencies (>12%), and had increased JH6 usage (63.9% JH6 usage vs 19.1% in total memory cells; data not shown). This population is reminiscent of class switched IgD B cell populations, mostly observed in germinal centers, that are reported to have high mutation rates (>10%) and preferential usage of JH6 segments (Zheng et al. 2004; Koelsch et al. 2007; Budeus et al. 2023; Seifert and Küppers 2016).

The Immcantation (Gupta et al. 2015) suite of tools was used to call clones based on the heavy chain junction length and sequence identity; subclones were called by light chains when available. 1219 clones had multiple members with cluster sizes ranging from 2-16 (Fig. 5E). Certain clones varied in their heterogeneity; for example, a group containing 14 IgD clones had more divergent sequences than a more tightly clustered group of 11 IgA2 clones.

### Antibody synthesis and binding results

To investigate the function of novel antibody sequences identified in the ASCs, we generated 100 monoclonal antibodies based on consensus heavy and light chain sequences. The first batch of 50 antibodies were synthesized as full-length IgG1 immunoglobulins, and the subsequent 50 were engineered as single-chain variable fragment (scFv)-IgG1-Fc fusions. This approach allowed for both the assessment of native antibody binding characteristics (full-length IgG1) and the rapid screening of multiple antibody specificities simultaneously (see Methods).

Initially, the first batch of antibodies were screened for binding to measles, mumps, and rubella virus antigens via commercial ELISA kits. One of the full-length IgG1 antibodies, MMR-32 yielded positive binding results. MMR-32 displayed apparent polyreactivity as it bound to ELISA plates coated with both mumps and rubella virus antigens (Supplemental Fig. S8). None of these antibodies exhibited neutralizing activity against MeV, MuV or RubV (data not shown).

Subsequent screening of the 50 scFv-Fc fusions by FACS on ExpiCHO cells transiently transfected with MeV and MuV fusion (F) and receptor binding protein (RBP) expression constructs identified several clones demonstrating binding to MeV-F proteins (Supplemental Fig. S9), notably MMR-82 and MMR-205. Further characterization confirmed MMR-82 binding to native MeV envelope proteins on infected Raji-B cells (Fig. 5F). Importantly, MMR-82 effectively neutralized MeV infection (Fig. 5G).

None of the antibodies, including the scFv-Fc fusions, demonstrated positive binding to the viral antigens in this ELISA format while hyperimmune polyclonal sera demonstrated the expected reactivity in the ELISA assays (Supplemental Fig. S8, ScFV-Fc data not shown). The lack of reactivity among the monoclonal antibodies highlights the possibility of false-negative results with ELISA-based screening methods.

### Germline specific repertoire analysis of antibody sequences confirm B cell class and allows haplotype distinction

Our pipeline allowed us to unambiguously assign IgH and IgL (Fig. 6A, Supplemental Fig. S10 & S11) gene family usage to our full-length antibody sequences, revealing distinct gene segment repertoires in cells collected from the MBC and ASC gates. Similarly, IgK repertoires were assigned using the total IMGT-GENE reference database and revealed distinct usage of VH segments across the three different populations (Supplemental Fig. S10). The MBCs exhibited a diverse VH and JH segment repertoire, with preferential usage of certain VH gene families, VH3 and VH4 (Fig, 6A) and JH4 segments (Supplemental Fig S11) known to be over-represented in MBCs in healthy adults (Tian et al. 2007). In contrast, the plasmablasts and plasma cells displayed a more restricted VH segment repertoire compared to the memory population. Notably, certain VH segments showed preferential enrichment or reduction in plasma cells and plasmablasts compared to MBCs, suggesting selective biases for antigen-specific antibody-secreting clones (Fig. 6A). A subset of sequences had low V segment identity (<85% V identity, or >15% mutation rate) when queried against the closest germline allele (7-9% for MBC and plasmablasts, 20% for plasma cells); these antigen-experienced B cells have higher mutation rates than expected for these populations. While 11% (31/286) of these highly mutated clones were in the MBCs and 13% (2/15) in the plasma cells were in the aforementioned IgD only population, the majority of the sequences were on IgG and IgA bearing cells. An increase in the proportion of highly mutated B cells is to be expected in a healthy adult immune system, given the propensity of increased exposures to the same immunizing antigens from natural re-infections or repeated vaccinations possibly recalling such B cells into further maturation rounds; however, it is important to validate the predicted mutations in such sequences in the context of any error biases possible from a given sequencing or enhancement technique. Often these sequences are filtered out as “warning sequences” in some high throughput statistical analysis programs like Hi-V Quest analysis from IMGT, requiring a closer evaluation. When we evaluated these sequences separately for VH usage (Fig. 6A, light blue bars), we found biases in these highly mutated sequences that were specific for each population; in some cases, sole usage of certain segments were found only in one population over the others for these sequences, like VH1-2 segments that were only found in the plasma cell population. This specific pattern of expression further highlights characteristics of B cell maturation through selection over any ubiquitous mutation error of any technique.

**Figure 6.**
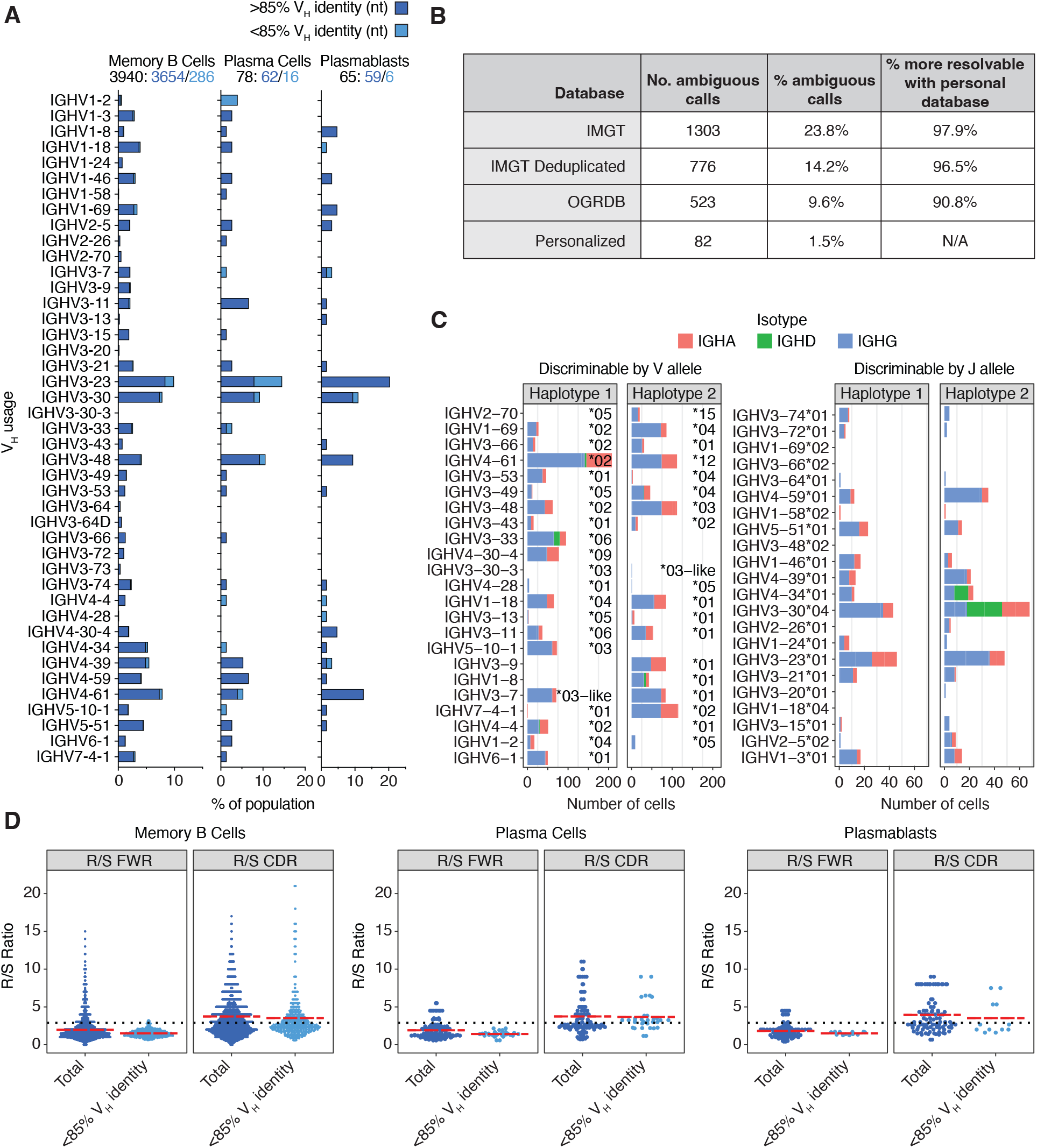
Donor-specific V-gene usage and mutation rates. (*A*) Overall IGHV gene usage frequency for all complete and productive heavy chain consensus sequences with an in-frame CDR3. Cells with a low V gene identity, high mutation rates (≥15%) are stratified (light blue), as they are sometimes filtered out by programs such as Hi-V-QUEST (IMGT). (*B*) Statistics for ambiguous IGH and IGL V gene calls when using IgBLAST with various databases. (*C*) V gene usage for cells expressing one of the heterozygous V alleles (left), or that expressed a homozygous V allele but were haplotype-discriminable by a heterozygous J allele (right). (*D*) Ratio of replacement to silent mutations (R/S) in FWR vs. CDR regions. The dotted line represents expected background R/S ratios for CDRS at R/S=2.9. The red lines are the mean R/S values. The elevated replacement mutations in the CDRs suggest cells that have undergone antigen selection.

When using a deduplicated IMGT-GENE database with IgBLAST, 14% of all functional V allele calls for IGH and IGL are ambiguously assigned to multiple equally likely alleles. When using a personalized reference database, ambiguous calls are reduced by over 90%; only 1.5% of calls remained ambiguous (Fig. 6B). The unambiguous calls are also needed to distinguish transcripts from each haplotype.

Sequences can be traced back to a specific haplotype if they include a gene or allele that is only present on one haplotype. 55% of IGH consensus sequences could be assigned a haplotype based on a heterozygous IGHV or IGHJ allele. 27% and 28% of consensus sequences came from haplotype 1 and 2, respectively. Only 0.3% of the consensus sequences had a contradictory haplotype assignment between the V allele and the J allele, which supports the phasing accuracy of the assembly and is likely the result of hypermutation. Although the overall usage of both haplotypes is similar, certain alleles are used more often than others. For example, IGHV7-4-1 was present on both haplotypes, but the *02 allele was used by 115 cells, compared to only a single cell using the *01 allele (Fig. 6C). The latter is known to be a poorly expressed VH allele in humans (Ohlin 2020). Two other poorly expressed human IGHV alleles (IGHV1-2*05 and IGHV4-4*01), known to be in linkage disequilibrium and typically found on the same haplotype (Ohlin 2020), were also present in our haplotype 2 and were found in 9 and 0 cells, respectively (Fig. 6C). Altogether, these data support the veracity of our analytic pipeline.

Mutational analysis of IGH shows an elevated ratio of replacement to silent mutations (R/S) in the CDR regions (mean = 3.7 - 3.9 depending on cell type) compared to the framework regions (FWR) and to the rate for random mutation for CDRs (2.9) (Fig. 6D). This is indicative of B cell populations that have undergone antigen-dependent maturation and selection in germinal center reactions (Wrammert et al. 2008; Dörner et al. 1997). The R/S pattern holds true for the consensus sequences with lower IGHV nucleotide identity, supporting their authenticity.

## Discussion

Our study shows that nanopore full-length, single-cell transcriptome sequencing can be used to explore multiple important features of the reactive antibody repertoire. We found that active antibody secreting cells like the plasmablasts and plasma cells found in the CD27++/CD38++ ASC gate expressed a sufficient number of immunoglobulin heavy and light chain transcripts to enable the creation of high-quality, full-length consensus sequences without the need for any enrichment approaches that would bias the expression levels observed in each cell. Importantly, these consensus sequences captured not just the V(D)J domains, but also the complete C gene segments (and thus the antibody isotype), as well as the 3’ M exons that determine whether the antibody is destined to be secreted or trafficked to the cell membrane. Unambiguous isotype assignments are crucial for understanding the downstream effector function of each antibody and its role in the larger response to an immune challenge.

As expected, expression of heavy and light chain transcripts was lower in the memory B cell population, requiring a library enrichment step to obtain consensus sequences. The consensus sequences obtained from the MBCs revealed an antigen-experienced repertoire that reflected the diversity of single antibodies and only a modest number of clonal expansions. By sequencing full-length IG transcripts from thousands of individual MBCs, we were able to survey the landscape of antibody features in the population, including isotype, V gene usage, CDR3 lengths, and mutation rates. We observed V gene usage frequencies in our heavy and light chain cohorts commonly seen in repertoires of post germinal center, isotype switched B cells (Glanville et al. 2009, 2011; Tian et al. 2007). These include higher frequencies of VH3 and VH4 families in the heavy chains, VK1 and VK3 families in the kappa chain, and VL1 families in the lambda chains. V gene repertoire was also influenced by selection, as we observed differing patterns of V segment usage between the memory and antibody-secreting populations, implying positive selection of antibody-producing B cells with favorable antigen binding attributes.

Sequencing of the full-length IG transcripts permitted a complete evaluation of the full spectrum of CDR3 lengths (between 4-37aa) in our collected antibody repertoire, whereas short read technologies might truncate especially long CDR3 sequences. While we observed a mean average CDR3 length in our MBC population that is standard in mature B cells (∼14aa), we were able to capture a rare few with CDR3 lengths over 30aa. While the function of this group is unknown without further evaluation, long CDR3 antibodies, while often rare due to increased autoreactive potential (Wardemann et al. 2003; Georgiou et al. 2014; Larimore et al. 2012), have significantly been associated with a variety of protective traits in infectious disease, including associations with highly potent, broadly neutralizing antibody responses developed in patients with chronic infections like HIV (Yu and Guan 2014). These findings should be considered when choosing a sequencing platform for future antibody repertoire studies.

A matching whole transcriptome library from the MBC gate revealed preferential isoform expression in certain clusters of MBCs, indicating a level of heterogeneity within the population that is hidden when viewed at the level of gene expression. While we highlight one example of the critical apoptosis regulator BCL2, future studies will undoubtedly reveal numerous associations and mechanisms driven by isoform-level, rather than gene-level, expression patterns in single B cells.

The differential expression of two key glycosylation pathway genes, B4GALT1 (IgG1) and FUT8 (IgA1/IgA2), within specific antibody isotypes of plasmablasts and plasma cells (Fig. 4D-E) unveils a layer of functional specialization that may define the fine-tuning of immune responses. For instance, B4GALT1 is instrumental in the addition of GlcNAc moieties to N-glycans, while FUT8 is responsible for adding fucose residues to the basal GlcNAc core, modifications known to increase and dampen ADCC activity, respectively (Nimmerjahn and Werner 2021; Wang and Ravetch 2019). Furthermore, in the thousands of MBCs profiled in the MBC gate, high MGAT3 and SLC35C1expression was not only restricted to a minority of clones but was also mutually exclusive. This is consistent with the opposing roles of MGAT3 and SLC35C1 in increasing and decreasing ADCC activity, respectively (Fig. 4D and Supplemental Fig. S5). To our knowledge, this is the first report to associate expression of effector function modifying N-glycan genes to B cell clones at a single-cell level. The nuanced interplay between these glycosylation patterns and specific antibody isotypes could be pivotal in orchestrating immune responses, where subtleties in effector function can have profound implications for pathogen neutralization, autoimmunity, and therapeutic antibody design (Archer et al. 2022; Wang 2019). Given the personalized nature of the immune response, our observations underscore the potential of leveraging single-cell sequencing technologies to dissect the molecular underpinnings of antibody function in the future of precision medicine.

The consensus heavy and light chain sequences generated were highly accurate, represented by all ∼100 antibody sequences that we tested, and were viable as they successfully produced as full-length IgG1 or scFv-Fc fusions. At least one was a high-affinity binder and neutralizer for measles virus (MMR-82 in Fig. 5F,G), while several more were either low-affinity binders or polyreactive clones (MMR-32, MMR-205) that also bound to native cell surface antigens. In the absence of antigen baiting, it is unclear what the “hit-rate” would be for a MMR vaccine since this has not been attempted before. Additionally, the wide diversity of apparently functional clones without known cognate antigens constitutes the remaining structure-function relationships which are still not well understood for circulating antibody sequences.

Our focus in this study was to describe the antibody repertoire using a combination of single-cell, full-length transcriptomics and a haplotyped long-read assembly of the IGH and IGL loci. Bulk adaptive immune receptor repertoire (AIRR-seq (Ford et al. 2023)) data has been paired with matching IGH genome assemblies before, but those genomes were generated from targeted sequencing approaches, which result in shorter reads and more discontiguous assemblies (mean # of contigs > 100) (Rodriguez et al. 2023). In contrast, our donor’s IGH locus was assembled into a single contig and phase block, showing that highly contiguous and fully-phased IGH assemblies can be created and reliably annotated from standard long-read whole genome sequencing data.

We were able to successfully map the antibody repertoire from the plasmablasts, plasma cells, and MBCs to their cognate germline sequences, often in a haplotype-specific fashion where heterozygous V or J alleles permitted, with less V gene allele ambiguity than if relying on public databases for IgBLAST (1.5% ambiguous calls versus 9.6% - 23.8%). Our comprehensive list of alleles in both haplotypes reveals some that rarely or never produce productive transcripts, such as IGHV2-70*05.

Strategies similar to what has been described in this work may be useful for large-scale immunogenomic population studies and pangenome references covering a broad spectrum of IG allelic diversity. Larger datasets where single-cell B cell sequencing is paired with germline assembly will be critical for understanding haplotype-specific expression patterns for specific alleles, allelic silencing and imbalances, somatic hypermutation, and heavy-light chain pairing within individual B cells during cellular development.

## Methods

### Vaccination and study design

A combination measles, mumps, and rubella (MMR) vaccine (M-M-R II®, Merck and Co.) was administered subcutaneously to a consented donor. 6 days post-vaccination, 60 ml of whole blood was collected directly into BD Vacutainer CPT tubes (Becton Dickinson) and PBMCs were isolated by centrifugation at 1,500x g for 20 minutes at room temperature according to the manufacturer’s instructions. The buffy coat containing PBMCs were removed by pipet, washed twice with 1X PBS and counted by hemocytometer before further processing for B cell enrichment.

### B cell enrichment and FACS

B cells were isolated using the Miltenyi B cell Isolation Kit II for negative B cell enrichment (Miltenyi Biotech, Auburn, CA) and passed over a Miltenyi LS column on a QuadraMACS magnetic separator with FACS buffer (1X PBS, 0.5% BSA, and 2 mM EDTA) according to manufacturer instructions. The collected flow-through, which contained untouched, enriched B cells, was washed twice in FACS Buffer and counted by hemocytometer with trypan blue staining (0.4% Trypan Blue in PBS solution) at a 1:10 dilution. The non-B cell fragment magnetically adsorbed to the column was also collected by removing the column from the magnet and applying pressure with the included plunger. The non-B cell fragment was collected in a separate 15 ml conical, counted and resuspended in FACS buffer.

Enriched B cells were resuspended in ice cold FACs buffer at a density of 10^6^ cells/100 μL and treated with Human Fc Block (BD) according to manufacturer instructions. After blocking, the fraction was stained with an antibody-secreting/memory B cell specific cocktail for FACS: Ovalbumin-AF488 (ThermoFisher), mouse anti-human IgM-AF488 (clone SA-DA4, SouthernBiotech, Birmingham, AL), mouse anti-human CD3-FITC (clone 7D6, ThermoFisher), mouse anti-human CD4-AF488 (clone RFT4, SouthernBiotech), mouse anti-human CD8-AF488 (clone RFT8, SouthernBiotech), mouse anti-human CD16-AF488 (clone 3G8, SouthernBiotech), mouse anti-human CD19-BV421 (clone HIB19, BD), mouse anti-human CD27-PE (clone O323, BioLegend), and mouse anti-human CD38-AF647 (clone HIT2, BioLegend). Cells were treated for 1 hour in the dark on ice. After staining, cells were washed with 1X PBS (no BSA), resuspended in 1 ml 1X PBS and treated with LIVE/DEAD™ Fixable Near-IR Dead Cell Staining reagents (ThermoFisher) according to manufacture instruction. After live/dead staining cells were washed 3x in FACS buffer and resuspended in 200 μL FACS buffer for acquisition on a 5 laser BD FACS Aria II in the Icahn School of Medicine Flow Cytometry Shared Resource Facility. Right before sorter acquisition, cells were strained through a 35 μm snap strainer in a Falcon™ round bottom polystyrene tubes for sorting (ThermoFisher) to dislodge any large cell-cell adhesion events. B cells were separated from non-B cells by gating on the AF488/FITC negative gate against the FITC positive “non-B cell dump gate” separating CD3,4 and 8 T cells and CD16 NK cells and monocytes. B cells were further enriched by CD19+ gating and subgated for CD27 and CD38 expression. Antibody-secreting cells (ASC gate, CD27++/CD38++) were separated from resting B memory cells (CD27+/CD38-) based on CD27 and CD38 expression. Antibody-secreting and resting memory B cells were collected in sterile Falcon tubes as the sorted populations.

### DNA extraction and germline whole genome sequencing

Genomic DNA was extracted from the monocyte cellular fraction using the Puregene Cell Kit (Qiagen) according to the modified Oxford Nanopore Technologies protocol available here: https://community.nanoporetech.com/extraction_methods/cell-line-dna-pure. Extracted DNA was further purified by isopropanol precipitation. DNA was quantified using a Qubit fluorometer (Thermo Fisher). Extracted genomic DNA was prepared for whole genome sequencing following the Ligation Sequencing Kit protocol (Oxford Nanopore Technologies, catalog no. SQK-LSK114) with long fragment buffer. The resulting libraries were sequenced on a PromethION (Oxford Nanopore Technologies) with R10.4.1 flow cells (Supplemental Table S3).

### Germline immune repertoire annotation

WGS reads were assembled into a haploid assembly using *Flye* (Kolmogorov et al. 2019) (--nano-hq). The assembly was converted into a diploid assembly using the *HapDup* (Kolmogorov et al. 2023) pipeline. Contigs were mapped against the GRCh38 human reference sequence with *minimap2* (Li 2018) (-ax asm10) to determine which contigs corresponded to IG regions. To annotate the assembly, V and C alleles from the IMGT GENE-DB (Li 2018; Giudicelli et al. 2005) and reference sequences were queried against the assemblies using *BLASTN* (Camacho et al. 2009) with “-penalty −5”. D and J alleles were queried with “-penalty −4 -task blastn-short”. In each locus, the allele with the highest alignment score (AS) was selected. Loci where the best alignment is not 100% identical to the existing best hit are potentially novel alleles; these alleles were validated with manual inspection of the raw-read alignments. Custom blast databases for use with *IgBLAST* (Ye et al. 2013) were created from the *seqkit* deduplicated (Shen et al. 2016) IGH and IGL V, D, and J alleles that were extracted from the assembly, along with reference C genes and IGK alleles from the IMGT GENE-DB.

### 10X Chromium processing

Sorted B cells in the ASC and MBC gates were separately processed for single cell RNAseq using the Chromium Next GEM Single cell 3’ Reagent kits v3.1 (10X Genomics) according to the manufacturer instructions with the following modifications. After the full-length cDNA amplification step post GEM-RT cleanup, 50% of the cDNA samples from each sorted cell population were saved for long read cDNA sequencing by Oxford Nanopore Technologies. The other 50% were processed for 3’ gene expression library construction, subjected to size selection by SPRIselect reagent beads, quality controlled for library construction via Agilent Bioanalyzer before Illumina sequencing on a HiSeq2000.

### Long read single cell transcriptome sequencing

10X cDNA was re-amplified using forward (10x_cDNA_fwd, read1) and reverse primers (10x_cDNA_rev, TSO) (Supplemental Table S4). Amplification reactions consisted of 50 μL LongAmp Taq 2x Master Mix (New England Biolabs), 1 μL 10 μM forward primer, 1 μL 10 μM reverse primer and 4 ng cDNA input in a final volume of 100 μL. Thermal cycling was performed using 30 seconds at 95°C for initial denaturation, followed by 14 cycles of 15 seconds at 95°C for denaturation, 15 seconds at 60°C as annealing, 6 minutes at 65°C for extension, and final extension at 65°C for 10 minutes. PCR products were purified using 0.7x v/v AMPure XP (Beckman Coulter) with 80% ethanol washes and eluted into 10 μL of water for downstream processing. DNA was quantified using the Qubit fluorometer (Thermo Fisher). The ASCs (ASC gate) were prepared for sequencing following the standard Ligation Sequencing Kit (Oxford Nanopore Technologies, catalog no. SQK-LSK110) with short fragment buffer using 200 ng input of amplified cDNA. The resulting single-cell, whole transcriptome libraries were sequenced on a PromethION (Oxford Nanopore Technologies) with R9.4.1 flow cells. The MBC gate library was prepared for sequencing following the single-cell cDNA protocol (Oxford Nanopore Technologies, catalog no. SQK-PCS111) using 10 ng input of enriched cDNA. The resulting single-cell, whole transcriptome libraries were sequenced on a PromethION (Oxford Nanopore Technologies) with R9.4.1 flow cells.

### IG transcript-enriched transcriptome sequencing

10X cDNA was re-amplified as described above. To increase IG heavy and light chain transcript coverage, an xGen™ Custom Hybridization Panel (Integrated DNA Technologies) was designed targeting IG and TCR exons. The hybridization panel consisted of 1877 5’ biotinylated probes, pooled at equimolar concentrations (Supplemental Table S5). The sample from the MBC gate was prepared starting with 500 ng of amplified 10x cDNA library. The manufacturer protocol was followed with the following modifications: no blocking oligos were used and Dynabeads kilobaseBINDER M-280 Streptavidin beads were used instead of Dynabeads M270 Streptavidin beads. Additionally, the post-capture PCR of the sample-bead conjugate was performed with 25 μL LongAmp Taq 2x Hot Start Master Mix (New England Biolabs, M0533), 1 μL 10 μM forward (10x_cDNA_fwd, read 1) primer, and 1 μL 10 μM reverse (10x_cDNA_rev, TSO) primer (Supplemental Table S4) following the long amplification PCR protocol. Thermal cycling was performed using 30 seconds at 95°C for initial denaturation, followed by 14 cycles of 15 seconds at 95°C for denaturation, 15 seconds at 60°C as annealing, 6 minutes at 65°C for extension, and final extension at 65°C for 10 minutes. PCR products were purified using 1.5x v/v AMPure XP (Beckman Coulter) with 80% ethanol washes and eluted into 25 μL of water for downstream processing. DNA was quantified using the Qubit fluorometer (Thermo Fisher). After post-capture PCR, only 300 ng of the cDNA sequencing library was prepared following the Ligation Sequencing Kit (Oxford Nanopore Technologies, catalog no. SQK-LSK110) with short fragment buffer. The resulting single-cell, IG transcript-enriched libraries were sequenced on a PromethION (Oxford Nanopore Technologies) with R9.4.1 flow cells.

### Expression analysis

Illumina reads were processed using the *cellranger count* command from the Cellranger software package using default parameters to generate single-cell gene expression count matrices. Single-cell gene and transcript expression count matrices from the Oxford Nanopore reads were generated using the *wf-single-cell pipeline* (https://github.com/epi2me-labs/wf-single-cell). The gene expression count matrices were imported into Seurat v5 (Hao et al. 2024) using the command *CreateSeuratObject* with the following parameters: min.cells = 3 and either min.features = 200 (for Illumina reads from MBC gate) or min.features = 100 (for the Oxford Nanopore reads from the ASC gate). Gene expression matrices were further filtered for both Illumina and Oxford Nanopore data in Seurat to retain only cell barcodes with nFeature_RNA values between 1500 and 9000. The transcript expression matrix from the Oxford Nanopore sequencing was similarly imported into Seurat using min.cells = 3 and min.features = 200, then filtered to only retain cell barcodes with nFeature_RNA values between 200 and 7500. Finally, all expression count matrices were scaled, normalized to 10,000 counts per cell barcode, and log transformed using the Seurat command *NormalizeData*.

### Doublet analysis

Doublets were identified by *scDblFinder* (Germain et al. 2021) with default settings, using Cellranger’s filtered_feature_bc_matrix for the Illumina data or *wf-single-cell*’s gene_expression.counts.tsv for the Oxford Nanopore data. Because doublets are difficult to detect among low complexity cell populations (e.g. homotypic doublets), cell barcodes were considered doublets and removed from subsequent analysis if they were detected in either the Oxford Nanopore or Illumina sequencing data.

### Antibody consensus sequences

Strand-oriented and full-length (i.e. bounded by the expected 10X read1 and TSO adapter sequences) reads were identified as part of the *wf-single-cell* pipeline. Those reads aligning to the IGH, IGK, and IGL loci were then identified by querying all reads with IgBLAST (Ye et al. 2013) using the personalized database and parameters “-c_region_db ncbi_human_c_genes - auxiliary_data human_gl.aux -outfmt 19”. Non-chimeric IG reads were then filtered based on the following criteria, keeping only reads where: (1) e-values for the annotated V and C segments were <1e-6, (2) annotated CDR3 regions were between 3-50 amino acids long, (3) alignments of the V and C segments covered >90% of the reference sequence (secreted form for IGHC), (4) annotated C and V segments were separated by <250 bp, and (5) the read contained <400 bp before the V segment or after the C segment.

Reads surviving this filtering were separated based on their IGK, IGL, or IGH locus assignment and associated with their cell barcode assignment from the *wf-single-cell* pipeline. Reads were downsampled where necessary to no more than 150 reads per cell barcode, per locus. Consensus sequences for each cell barcode and locus were generated using *medaka* (https://github.com/nanoporetech/medaka) with the *smolecule* subcommand, using the parameters “--min_depth 3 --depth 10 --model r941_e81_sup_g514”. Finally, the consensus sequences were queried with *IgBLAST* using existing reference databases (IMGT GENE-DB (Ye et al. 2013; Giudicelli et al. 2005), OGRDB (Lees et al. 2020)) or a custom BLAST database built using the personalized assembly of IG regions, using the reference C and IGK sequences but replacing the V/D/J reference FASTAs with IGH/IGL fasta sequences extracted from the assembly (see Germline immune repertoire annotation). If multiple light chains were present in one cell, the light chain with the highest coverage was used for subsequent analysis. Consensus sequences were filtered out if they were unproductive, had an incomplete VDJ, or were missing the conserved junction anchors (C104 and W/F116).

### Clonotype and Mutational Analysis

The *Immcantation* (Gupta et al. 2015) suite was used to call clones and do mutational analysis. Nearest neighbor distances were determined with the HH_S5F model (Yaari et al. 2013) and thresholds for distinguishing clones were determined using the “gamma-norm” curve. Clones were called with *SCOPer* (Nouri and Kleinstein 2020), using additional light chain information where available. Mutation profiling was done with *SHazaM* (Gupta et al. 2015).

IG network creation was done using ‘generate_network’ from *Dandelion* (Suo et al. 2024), which draws the network based on the Levenshtein distance between amino acid VDJ sequences.

### Antibody synthesis

The monoclonal antibodies derived from the sequences of unique barcoded ASC-gated B cells were synthesized and produced in two batches. For the first batch, 45 unique matched heavy and light variable sequences were chosen randomly from the total sorted ASC gate (214 cells in total). Variable sequences were synthesized and cloned by a CRO (GenScript, Piscataway, NJ) into in-house pcDNA3.4-based vectors containing a human G1 constant region for the heavy chain, a human kappa constant region for kappa sequences, and a human lambda constant 2 region for lambda sequences. Cloning was done in-frame of a vector supplied signal sequence and kozak;heavy and light chain vectors were co-transfected into Expi293 cells, and supernatants collected a week later (GenScript). Supernatants were quantitated for antibody concentrations by bio-layer interferometry (BLI) on an Octet Red 96 (Sartorius) using protein A biosensors and a human IgG1 isotype control antibody as a standard. For the second batch, 50 unique sequences were selected from the remaining ASC cohort, and heavy and light chain variable sequences were synthesized as single chain variable fragments (ScFvs) containing a interchain (Light-Heavy) G_4_S^3^ linker and cloned into a pFuse-hG1-Fc2 Fc-tagged vector (InVivoGen, San Diego, CA), in-frame between the vector supplied signal sequence (from the human Il-2 protein) and a human IgG1 Fc tag (Hinge+CH2+CH3 domains). Synthesis, cloning and vector DNA preps were done by CRO (Twist Biosciences, San Francisco, CA). Prepped pFuse ScFv-hG1Fc vectors (3 μg) were transfected into 7.5 x 10^6^ Expi293 cells with Expifectamine transfection reagent (ThermoFischer) according to manufacture instruction and grown in Expi293 serum free media (ThermoFisher) in 6 well tissue culture plates (3 ml total volume). After 5 days, supernatants were collected and quantitated by BLI on an Octet Red 96 (Sartorius) using protein A biosensors.

### Neutralization assays

Vero-CCL81 [for MuV and VSV_RUBV] or Vero-hSLAM [for MeV-IC323] target cells which were seeded at a density of 2 × 10^4^ cells per well in flat-bottom 96-well plates (Fisher Scientific, #08-772-3). The cells were incubated at 37 °C/5% CO_2_ overnight (∼20 hours). On the following day, GFP-expressing vaccine strain MuV-JL5, recombinant MeV-IC323, or recombinant VSV virions expressing Rubella E2/E1 envelope region were pre-incubated with serially diluted antibodies for a minimum of 30 minutes at 37°C (MuV and MeV-IC323) or for 1 hour at 32°C (for VSV_RUBV), and then added to the target cells. Between 16-24 hours post-infection, GFP counts were acquired by the Celigo imaging cytometer (Nexcelom Biosciences, version 4.1.3.0)

### ELISA

ELISA kits against Measles, Mumps and Rubella were purchased commercially (Arlington Scientific, # 850096AG, # 860096AG and # 800096AG) and performed according to manufacturer’s instructions. Antigen substrates were sonicated infected cell lysates. Briefly, the antibody samples and the various controls were first diluted, then added to the wells. The plates were then incubated at room temperature for 30 minutes, washed 4X, after which HRP-conjugated secondary anti-human antibodies were added and the plates incubated for another 30 minutes at room temperature. Wells were then washed 4 more times; HRP substrate added and incubated an additional 30 minutes at room temperature. Stop reagent was added and absorbance was read by the CYTATION Gen5 Microplate reader (BioTek, part of Agilent Technologies, version 1.15) at 405 nm against reagent blank.

### Cell surface binding assays

Antibody binding to cell surface expressed mumps and measles envelope glycoproteins were performed two different ways. For initial screening, pCAGG expression plasmids containing MuV-F, MuV-RBP/HN, MeV-F, and MeV-RBP/H under the CMV enhancer and chicken beta actin promoter were independently transfected into Expi293 cells and then processed for FACS analysis 48 hours post-transfection. Cells were washed 3x in FACS Buffer (1X PBS, 0.5% BSA, 2 mM EDTA) and resuspended at a density of 1 x 10^6^ cells/ml plated (50 μL) in individual wells of a 96 well plate. Transfected supernatants from the synthesized recombinant monoclonal antibodies were then added 1:1 to the cells and stained for 1 hour on ice. Cells were then washed 3x in FACS Buffer and stained on ice for 1 hour with goat anti-Human IgG Fc Alexa Fluor™ 647 secondary antibody (ThermoFisher), protected from light. Cells were washed again 3x and resuspended in 200 μL. Binding was read by flow cytometry on an IntelliCyte High Throughput Flow Cytometry (IntelliCyte/Sartorius).

For confirmation of putative positive MeV binders, antibodies were used to detect MeV envelope glycoproteins (F or RBP/H) expressed on infected Raji-DC-SIGN B cells. Briefly, RAji-DC-SIGN cells were infected at an MOI of 0.01 and after 3-5 days of infection, the cells were seeded at 5 × 10^4^ cells per well in U-bottom 96-well plates (Fisher Scientific; #163320). Two dilutions of antibody, 10 μg or 1 μg and 1:300 or 1:3000 dilution of control MMR-vaccinated sera were made up to a total volume of 100 μL and incubated with infected cells for 2 hours at 37 °C, then washed twice with FACS Buffer (1% BSA, 0.1% Sodium Azide, and 1 mM EDTA in PBS). After the 2nd wash, 1:2000 dilution of secondary Alexa FLUOR-647 GOAT-Anti human IgG [H+L] (Invitrogen; #A-21445) was added to the cells for 30 minutes followed by 3 washes with FACS Buffer, and the cells were fixed for 10 minutes in 2% Formaldehyde at room temperature before detection on the Attune NxT Flow Cytometer (ThermoFisher Scientific). FCS files were analyzed using FlowJo v10.

## Data access

The sequencing data generated in this study have been submitted to the NCBI BioProject database (https://www.ncbi.nlm.nih.gov/bioproject/) under accession number PRJNA1087553. Scripts used to generate consensus sequences are available on GitHub (https://github.com/LynnLy/ig_consensus_pipeline).

## Acknowledgements

B.L. acknowledges the Ward-Coleman estate for endowing the Ward-Coleman Chairs at the Icahn School of Medicine at Mount Sinai. This work is also funded in part by DoD CDMRP PR192188 and NIH R01 AI185102. C.S. acknowledges funding from NIH T32 AI07647. C.H. acknowledges funding from the Ministry of Science and Technology (MoST) post-doctoral fellowship.

## Author contributions

J.B., J.A.D., C.S., S.J., S.H., and B.L. conceived the project. S.H., S.J., and B.L. coordinated the project. J.B. coordinated the data analysis and was the lead writer. C.T., J.A.D., C.H., and S.K. prepared the samples and generated the data. J.B., L.L., J.A.D., and C.S. analyzed the data. L.L., A.S., and A.W.D. developed the analysis pipelines. J.B., L.L., J.A.D., C.T., C.S., D.J.T., S.H., and B.L. contributed to writing the manuscript.

## References

Archer EJ, Gonzalez JC, Ghosh D, Mellins ED, Wang TT. 2022. Harnessing IgG Fc glycosylation for clinical benefit. Curr Opin Immunol 77: 102231.

Boyd SD, Gaëta BA, Jackson KJ, Fire AZ, Marshall EL, Merker JD, Maniar JM, Zhang LN, Sahaf B, Jones CD, et al. 2010. Individual variation in the germline Ig gene repertoire inferred from variable region gene rearrangements. J Immunol 184: 6986–6992.

Budeus B, Kibler A, Küppers R. 2023. Human IgM-expressing memory B cells. Front Immunol 14: 1308378.

Camacho C, Coulouris G, Avagyan V, Ma N, Papadopoulos J, Bealer K, Madden TL. 2009. BLAST+: architecture and applications. BMC Bioinformatics 10: 421.

Di Noia JM, Neuberger MS. 2007. Molecular mechanisms of antibody somatic hypermutation. Annu Rev Biochem 76: 1–22.

Dong J, Cross RW, Doyle MP, Kose N, Mousa JJ, Annand EJ, Borisevich V, Agans KN, Sutton R, Nargi R, et al. 2020. Potent Henipavirus Neutralization by Antibodies Recognizing Diverse Sites on Hendra and Nipah Virus Receptor Binding Protein. Cell 183: 1536–1550.e17.

Dörner T, Brezinschek HP, Brezinschek RI, Foster SJ, Domiati-Saad R, Lipsky PE. 1997. Analysis of the frequency and pattern of somatic mutations within nonproductively rearranged human variable heavy chain genes. J Immunol 158: 2779–2789.

Dudley DD, Chaudhuri J, Bassing CH, Alt FW. 2005. Mechanism and control of V(D)J recombination versus class switch recombination: similarities and differences. Adv Immunol 86: 43–112.

Duty JA, Szodoray P, Zheng N-Y, Koelsch KA, Zhang Q, Swiatkowski M, Mathias M, Garman L, Helms C, Nakken B, et al. 2009. Functional anergy in a subpopulation of naive B cells from healthy humans that express autoreactive immunoglobulin receptors. J Exp Med 206: 139–151.

Engblom C, Thrane K, Lin Q, Andersson A, Toosi H, Chen X, Steiner E, Lu C, Mantovani G, Hagemann-Jensen M, et al. 2023. Spatial transcriptomics of B cell and T cell receptors reveals lymphocyte clonal dynamics. Science 382: eadf8486.

Ford EE, Tieri D, Rodriguez OL, Francoeur NJ, Soto J, Kos JT, Peres A, Gibson WS, Silver CA, Deikus G, et al. 2023. FLAIRR-Seq: A Method for Single-Molecule Resolution of Near Full-Length Antibody H Chain Repertoires. J Immunol 210: 1607–1619.

Georgiou G, Ippolito GC, Beausang J, Busse CE, Wardemann H, Quake SR. 2014. The promise and challenge of high-throughput sequencing of the antibody repertoire. Nat Biotechnol 32: 158–168.

Germain P-L, Lun A, Garcia Meixide C, Macnair W, Robinson MD. 2021. Doublet identification in single-cell sequencing data using. F1000Res 10: 979.

Giudicelli V, Chaume D, Lefranc M-P. 2005. IMGT/GENE-DB: a comprehensive database for human and mouse immunoglobulin and T cell receptor genes. Nucleic Acids Res 33: D256–61.

Giudicelli V, Duroux P, Rollin M, Aouinti S, Folch G, Jabado-Michaloud J, Lefranc M-P, Kossida S. 2022. IMGT Immunoinformatics Tools for Standardized V-DOMAIN Analysis. Methods Mol Biol 2453: 477–531.

Glanville J, Kuo TC, von Büdingen H-C, Guey L, Berka J, Sundar PD, Huerta G, Mehta GR, Oksenberg JR, Hauser SL, et al. 2011. Naive antibody gene-segment frequencies are heritable and unaltered by chronic lymphocyte ablation. Proc Natl Acad Sci U S A 108: 20066–20071.

Glanville J, Zhai W, Berka J, Telman D, Huerta G, Mehta GR, Ni I, Mei L, Sundar PD, Day GMR, et al. 2009. Precise determination of the diversity of a combinatorial antibody library gives insight into the human immunoglobulin repertoire. Proc Natl Acad Sci U S A 106: 20216–20221.

Gupta NT, Vander Heiden JA, Uduman M, Gadala-Maria D, Yaari G, Kleinstein SH. 2015. Change-O: a toolkit for analyzing large-scale B cell immunoglobulin repertoire sequencing data. Bioinformatics 31: 3356–3358.

Hansen MH, Cédile O, Abildgaard N, Nyvold CG. 2024. The potential of 3rd-generation nanopore sequencing for B-cell clonotyping in lymphoproliferative disorders. EJHaem 5: 290– 293.

Hao Y, Stuart T, Kowalski MH, Choudhary S, Hoffman P, Hartman A, Srivastava A, Molla G, Madad S, Fernandez-Granda C, et al. 2024. Dictionary learning for integrative, multimodal and scalable single-cell analysis. Nat Biotechnol 42: 293–304.

Irvine EB, Alter G. 2020. Understanding the role of antibody glycosylation through the lens of severe viral and bacterial diseases. Glycobiology 30: 241–253.

Jaffe DB, Shahi P, Adams BA, Chrisman AM, Finnegan PM, Raman N, Royall AE, Tsai F, Vollbrecht T, Reyes DS, et al. 2022. Functional antibodies exhibit light chain coherence. Nature 611: 352–357.

Kenter AL, Watson CT, Spille J-H. 2021. Igh Locus Polymorphism May Dictate Topological Chromatin Conformation and V Gene Usage in the Ig Repertoire. Front Immunol 12: 682589.

Kidd MJ, Jackson KJL, Boyd SD, Collins AM. 2016. DJ Pairing during VDJ Recombination Shows Positional Biases That Vary among Individuals with Differing IGHD Locus Immunogenotypes. J Immunol 196: 1158–1164.

Koelsch K, Zheng N-Y, Zhang Q, Duty A, Helms C, Mathias MD, Jared M, Smith K, Capra JD, Wilson PC. 2007. Mature B cells class switched to IgD are autoreactive in healthy individuals. J Clin Invest 117: 1558–1565.

Kolmogorov M, Billingsley KJ, Mastoras M, Meredith M, Monlong J, Lorig-Roach R, Asri M, Alvarez Jerez P, Malik L, Dewan R, et al. 2023. Scalable Nanopore sequencing of human genomes provides a comprehensive view of haplotype-resolved variation and methylation. Nat Methods 20: 1483–1492.

Kolmogorov M, Yuan J, Lin Y, Pevzner PA. 2019. Assembly of long, error-prone reads using repeat graphs. Nat Biotechnol 37: 540–546.

Larimore K, McCormick MW, Robins HS, Greenberg PD. 2012. Shaping of human germline IgH repertoires revealed by deep sequencing. J Immunol 189: 3221–3230.

Lees W, Busse CE, Corcoran M, Ohlin M, Scheepers C, Matsen FA, Yaari G, Watson CT, AIRR Community, Collins A, et al. 2020. OGRDB: a reference database of inferred immune receptor genes. Nucleic Acids Res 48: D964–D970.

Lefranc MP. 2001a. Nomenclature of the human immunoglobulin heavy (IGH) genes. Exp Clin Immunogenet 18: 100–116.

Lefranc MP. 2001b. Nomenclature of the human immunoglobulin kappa (IGK) genes. Exp Clin Immunogenet 18: 161–174.

Lefranc MP. 2001c. Nomenclature of the human immunoglobulin lambda (IGL) genes. Exp Clin Immunogenet 18: 242–254.

Lefranc M-P, Pommié C, Kaas Q, Duprat E, Bosc N, Guiraudou D, Jean C, Ruiz M, Da Piédade I, Rouard M, et al. 2005. IMGT unique numbering for immunoglobulin and T cell receptor constant domains and Ig superfamily C-like domains. Dev Comp Immunol 29: 185–203.

Li H. 2018. Minimap2: pairwise alignment for nucleotide sequences. Bioinformatics 34: 3094– 3100.

Nimmerjahn F, Werner A. 2021. Sweet Rules: Linking Glycosylation to Antibody Function. Experientia Suppl 112: 365–393.

Nouri N, Kleinstein SH. 2020. Somatic hypermutation analysis for improved identification of B cell clonal families from next-generation sequencing data. PLoS Comput Biol 16: e1007977.

Nurk S, Koren S, Rhie A, Rautiainen M, Bzikadze AV, Mikheenko A, Vollger MR, Altemose N, Uralsky L, Gershman A, et al. 2022. The complete sequence of a human genome. Science 376: 44–53.

Ohlin M. 2020. Poorly Expressed Alleles of Several Human Immunoglobulin Heavy Chain Variable Genes are Common in the Human Population. Front Immunol 11: 603980.

Papalexi E, Satija R. 2018. Single-cell RNA sequencing to explore immune cell heterogeneity. Nat Rev Immunol 18: 35–45.

Peres A, Lees WD, Rodriguez OL, Lee NY, Polak P, Hope R, Kedmi M, Collins AM, Ohlin M, Kleinstein SH, et al. 2023. IGHV allele similarity clustering improves genotype inference from adaptive immune receptor repertoire sequencing data. Nucleic Acids Res 51: e86.

Rodriguez OL, Gibson WS, Parks T, Emery M, Powell J, Strahl M, Deikus G, Auckland K, Eichler EE, Marasco WA, et al. 2020. A Novel Framework for Characterizing Genomic Haplotype Diversity in the Human Immunoglobulin Heavy Chain Locus. Front Immunol 11: 2136.

Rodriguez OL, Safonova Y, Silver CA, Shields K, Gibson WS, Kos JT, Tieri D, Ke H, Jackson KJL, Boyd SD, et al. 2023. Genetic variation in the immunoglobulin heavy chain locus shapes the human antibody repertoire. Nat Commun 14: 4419.

Seifert M, Küppers R. 2016. Human memory B cells. Leukemia 30: 2283–2292.

Shen W, Le S, Li Y, Hu F. 2016. SeqKit: A Cross-Platform and Ultrafast Toolkit for FASTA/Q File Manipulation. PLoS One 11: e0163962.

Shen X, Klaric L, Sharapov S, Mangino M, Ning Z, Wu D, Trbojevic-Akmacic I, Pucic-Bakovic M, Rudan I, Polašek O, et al. 2017. Multivariate discovery and replication of five novel loci associated with Immunoglobulin G N-glycosylation. Nat Commun 8: 447.

Singh M, Al-Eryani G, Carswell S, Ferguson JM, Blackburn J, Barton K, Roden D, Luciani F, Giang Phan T, Junankar S, et al. 2019. High-throughput targeted long-read single cell sequencing reveals the clonal and transcriptional landscape of lymphocytes. Nat Commun 10: 3120.

Subas Satish HP, Zeglinski K, Uren RT, Nutt SL, Ritchie ME, Gouil Q, Kluck RM. 2022. NAb-seq: an accurate, rapid, and cost-effective method for antibody long-read sequencing in hybridoma cell lines and single B cells. MAbs 14: 2106621.

Suo C, Polanski K, Dann E, Lindeboom RGH, Vilarrasa-Blasi R, Vento-Tormo R, Haniffa M, Meyer KB, Dratva LM, Tuong ZK, et al. 2024. Dandelion uses the single-cell adaptive immune receptor repertoire to explore lymphocyte developmental origins. Nat Biotechnol 42: 40–51.

Tian C, Luskin GK, Dischert KM, Higginbotham JN, Shepherd BE, Crowe JE Jr. 2007. Evidence for preferential Ig gene usage and differential TdT and exonuclease activities in human naïve and memory B cells. Mol Immunol 44: 2173–2183.

Tonegawa S. 1983. Somatic generation of antibody diversity. Nature 302: 575–581.

Wahl A, van den Akker E, Klaric L, Štambuk J, Benedetti E, Plomp R, Razdorov G, Trbojevic-Akmacic I, Deelen J, van Heemst D, et al. 2018. Genome-Wide Association Study on Immunoglobulin G Glycosylation Patterns. Front Immunol 9: 277.

Wang TT. 2019. IgG Fc Glycosylation in Human Immunity. Curr Top Microbiol Immunol 423: 63–75.

Wang TT, Ravetch JV. 2019. Functional diversification of IgGs through Fc glycosylation. J Clin Invest 129: 3492–3498.

Wardemann H, Yurasov S, Schaefer A, Young JW, Meffre E, Nussenzweig MC. 2003. Predominant autoantibody production by early human B cell precursors. Science 301: 1374– 1377.

Warren CFA, Wong-Brown MW, Bowden NA. 2019. BCL-2 family isoforms in apoptosis and cancer. Cell Death Dis 10: 177.

Watson CT, Glanville J, Marasco WA. 2017. The Individual and Population Genetics of Antibody Immunity. Trends Immunol 38: 459–470.

Watson CT, Steinberg KM, Huddleston J, Warren RL, Malig M, Schein J, Willsey AJ, Joy JB, Scott JK, Graves TA, et al. 2013. Complete haplotype sequence of the human immunoglobulin heavy-chain variable, diversity, and joining genes and characterization of allelic and copy-number variation. Am J Hum Genet 92: 530–546.

Wrammert J, Smith K, Miller J, Langley WA, Kokko K, Larsen C, Zheng N-Y, Mays I, Garman L, Helms C, et al. 2008. Rapid cloning of high-affinity human monoclonal antibodies against influenza virus. Nature 453: 667–671.

Yaari G, Vander Heiden JA, Uduman M, Gadala-Maria D, Gupta N, Stern JNH, O’Connor KC, Hafler DA, Laserson U, Vigneault F, et al. 2013. Models of somatic hypermutation targeting and substitution based on synonymous mutations from high-throughput immunoglobulin sequencing data. Front Immunol 4: 358.

Ye J, Ma N, Madden TL, Ostell JM. 2013. IgBLAST: an immunoglobulin variable domain sequence analysis tool. Nucleic Acids Res 41: W34–40.

Yu L, Guan Y. 2014. Immunologic Basis for Long HCDR3s in Broadly Neutralizing Antibodies Against HIV-1. Front Immunol 5: 250.

Zheng N-Y, Wilson K, Wang X, Boston A, Kolar G, Jackson SM, Liu Y-J, Pascual V, Capra JD, Wilson PC. 2004. Human immunoglobulin selection associated with class switch and possible tolerogenic origins for C delta class-switched B cells. J Clin Invest 113: 1188–1201.

GitHub - nanoporetech/medaka: Sequence correction provided by ONT Research. GitHub. https://github.com/nanoporetech/medaka (Accessed March 13, 2024).

