## Supplemental Materials for "De novo antibody discovery in human blood from full-length single B cell transcriptomics and matching haplotyped-resolved germline assemblies"

### Supplementary Materials

#### Figure S1: Novel IGLC Allele

IGV pileup of closest matching reference alleles for novel IGLC gene segment. The colored vertical bars are mismatches between the assembled contig and the reads or IGLC alleles. Top: Raw reads aligned to the assembly. Bottom: Known IGLC alleles aligned to the assembly. The closest known matches have at least 1 SNP difference from the assembly and the genomic reads.

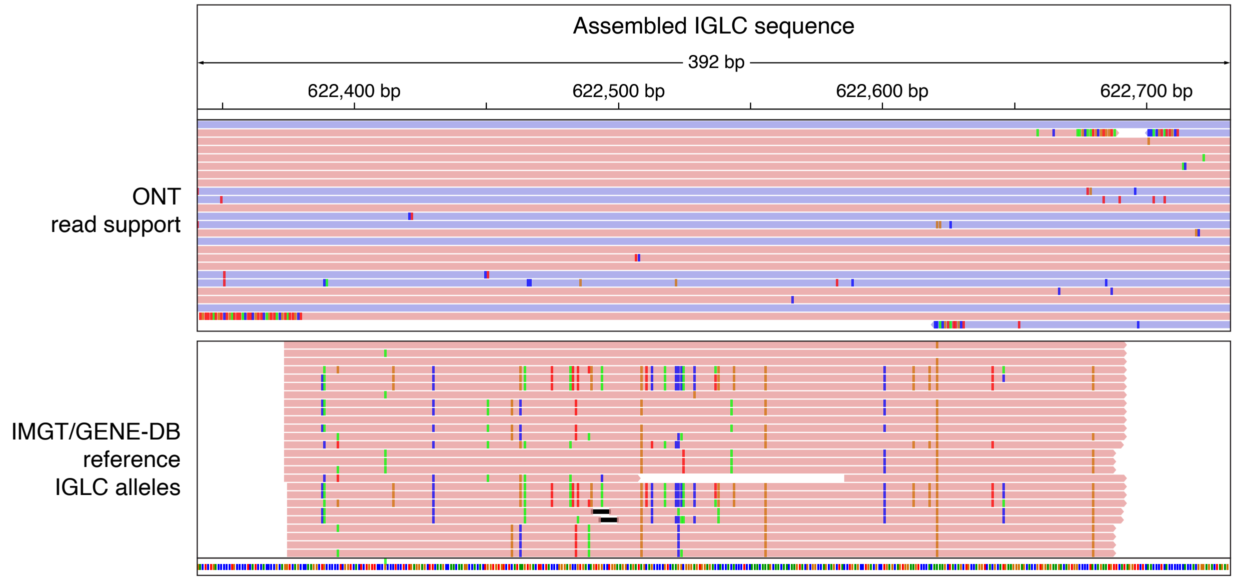

#### Figure S2: FACS plots

PBMCs were first enriched for B-cells via negative selection and sorted for memory B cells and antibody-secreting B cells according to the strategy shown. B cell enriched PBMCs were gated to retain only singlets and live cells (P4 gate). Cells in the P4 gate were then gated on CD19+ B cells that were IgM-. A cocktail of antibodies corresponding to T cells (CD3/CD4/CD8) and monocytes/NK cells (CD16) were further added to ensure purity of CD19+/IgM- B cells (P5 gate). P5 gate cells were sorted for antibody-secreting cells (CD38++/CD27++) and memory B cells (CD38-/CD27+) in the P7 gate (referred to as ASC gate) and P8 gate (referred to as MBC gate), respectively.

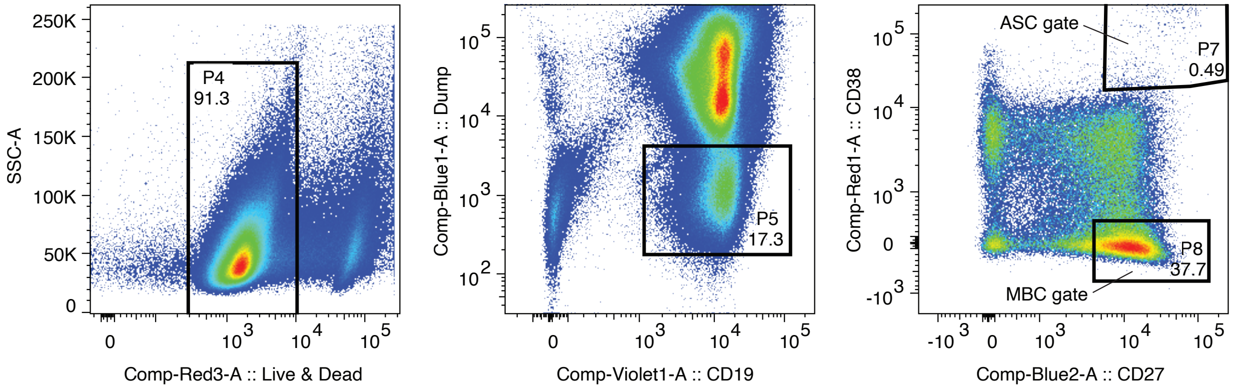

#### Figure S3: ASC gate UMI counts per cell barcode

UMI counts per cell barcode in Illumina and ONT for the single-cell library generated from the FACS P7 (ASC) gate.

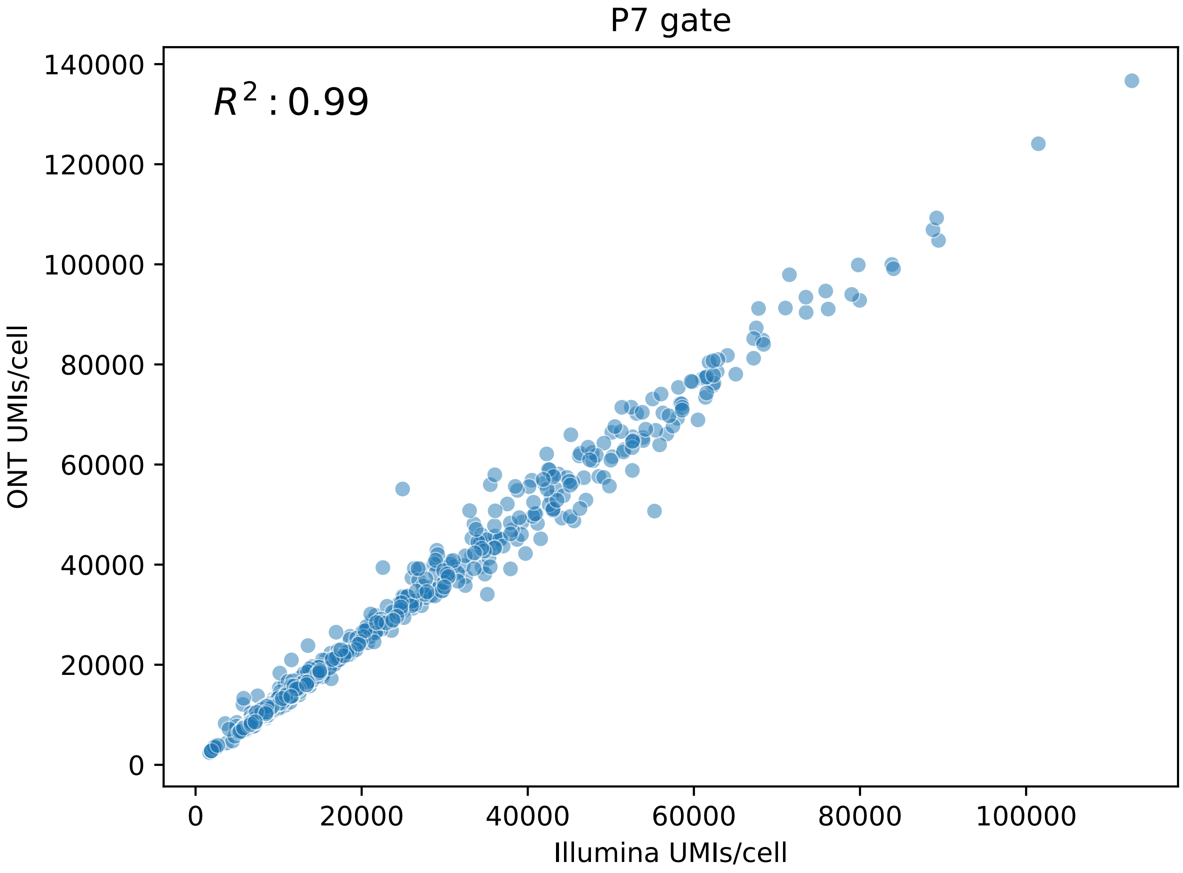

#### Figure S4: MBC gate UMI counts per cell barcode

UMI counts per cell barcode in Illumina and ONT for the single-cell library generated from the FACS P8 (MBC) gate.

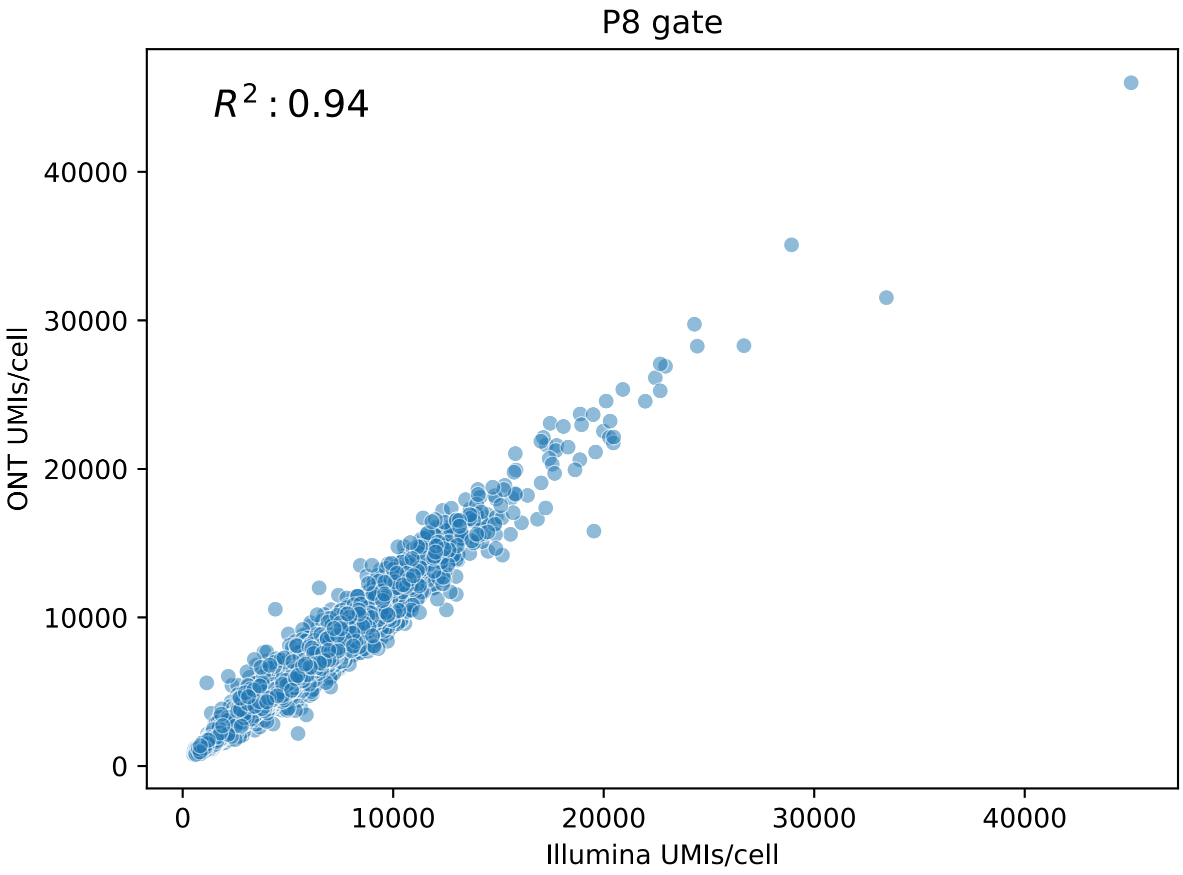

#### Figure S5: Glycosylation gene expression

Expression of glycosylation genes in memory B cells from the FACS MBC gate.

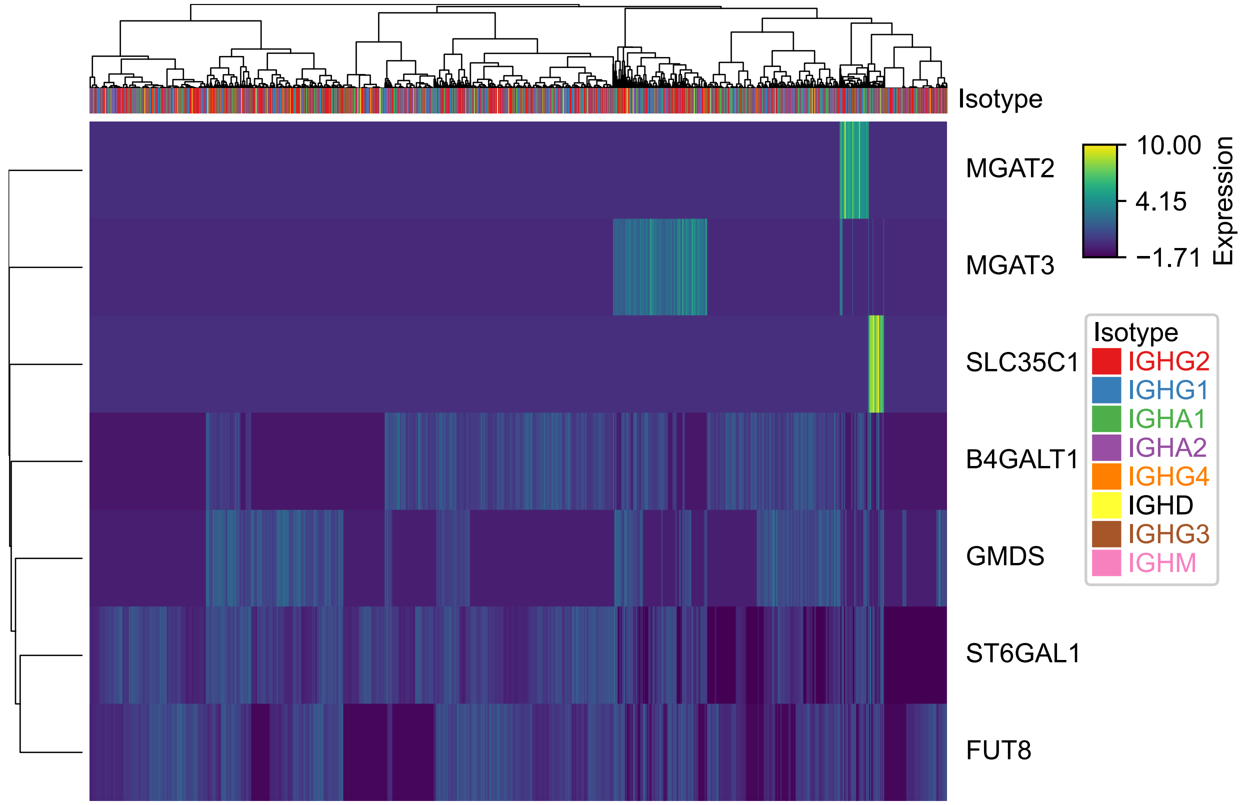

#### Figure S6: IG consensus pipeline

Pipeline used to create consensus heavy and light chain sequences for each cell barcode.

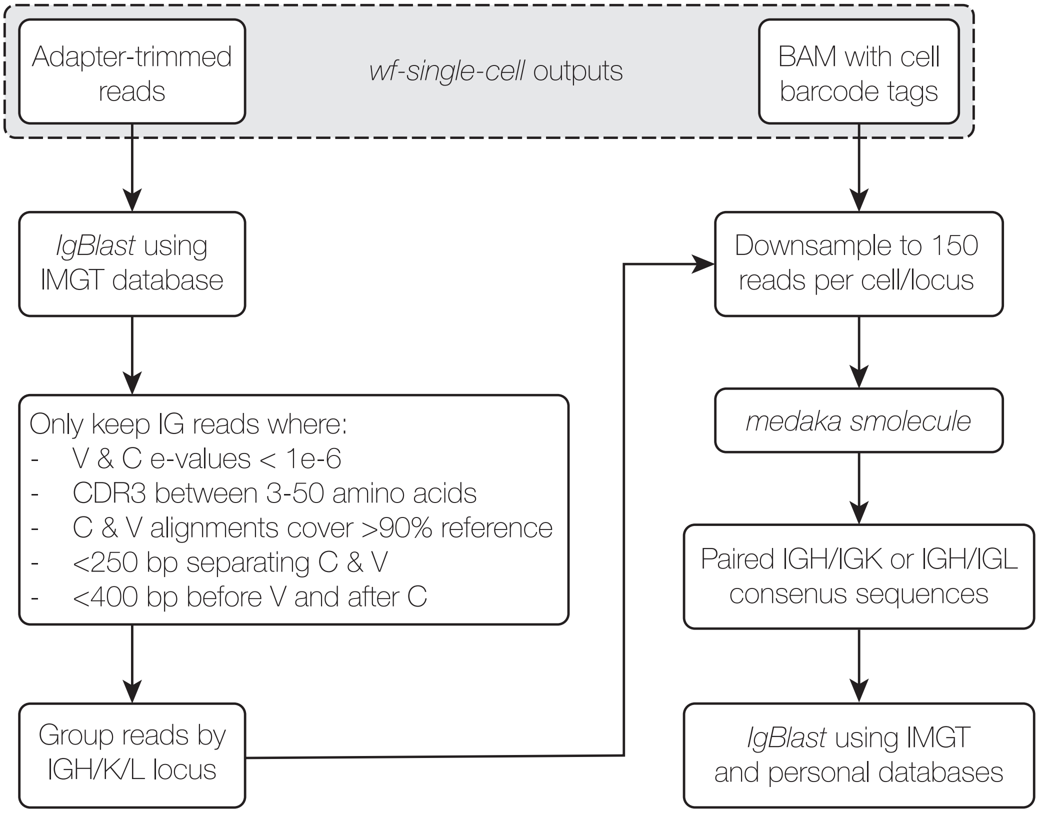

#### Figure S7: Isotype lengths

Nucleotide consensus sequence length distributions by locus (IGH, IGK, and IGL) and isotype.

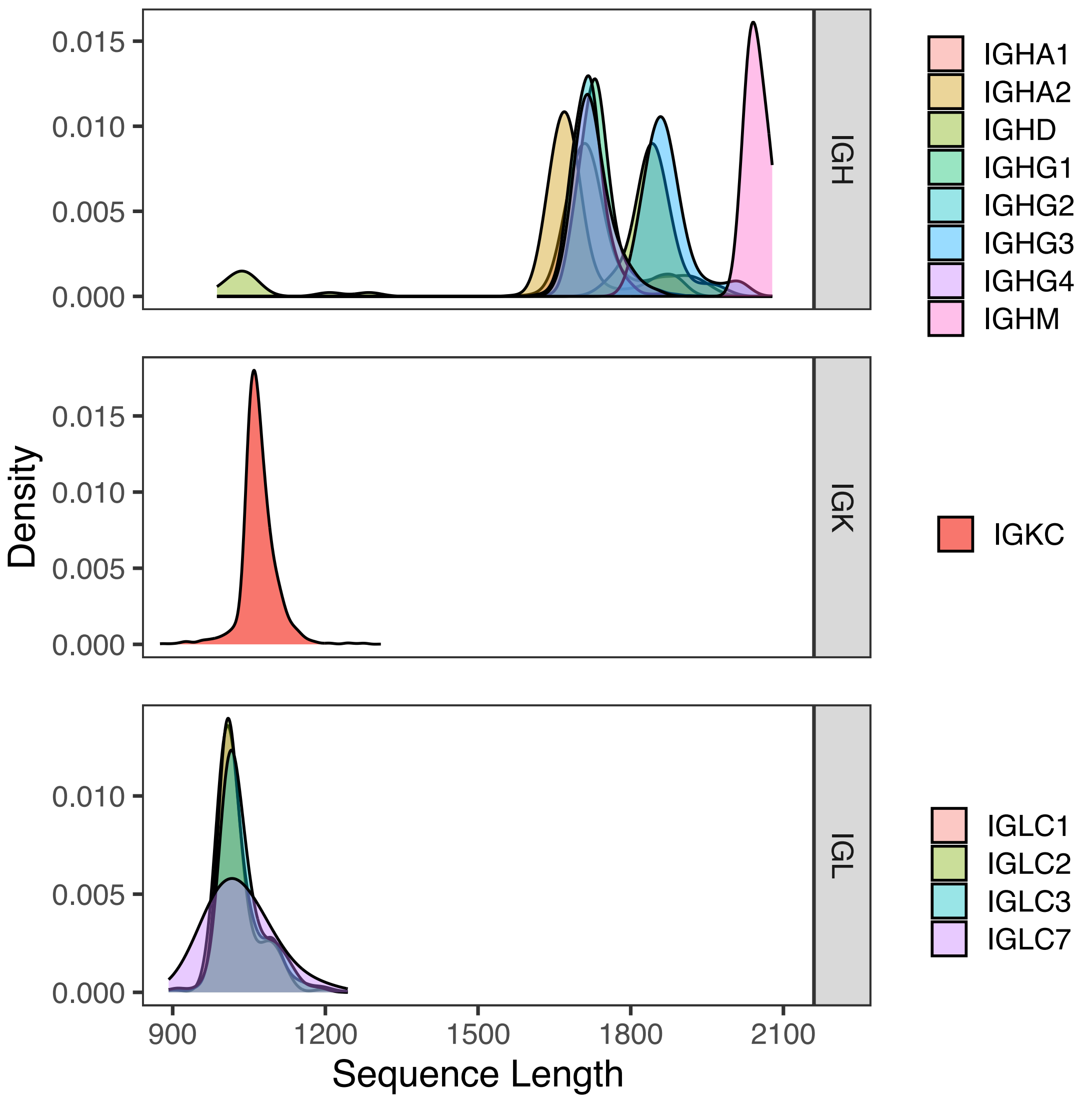

#### Figure S8: ELISA assay results

ELISA assay results.

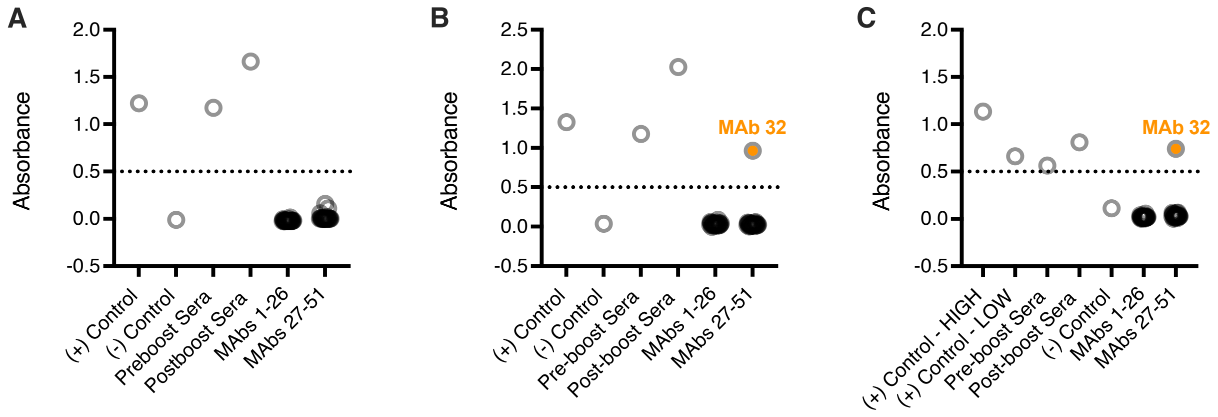

#### Figure S9: Binding assay results

ExpiCHO cells were transfected with pCAGG mammalian expression constructs containing MeV F, MeV RBP, MuV F or MuV HN proteins and stained with the ScFv-human G1-Fc monoclonal antibodies from our second batch of clones. Stained cells were detected with anti-human IgG-APC secondary and mean fluorescent intensities were recorded. Graph represents a select number of the clones, including the positive binders. Mock (empty plasmid) transfected cells and staining with no primary antibody (No Ab) were used as negative controls.

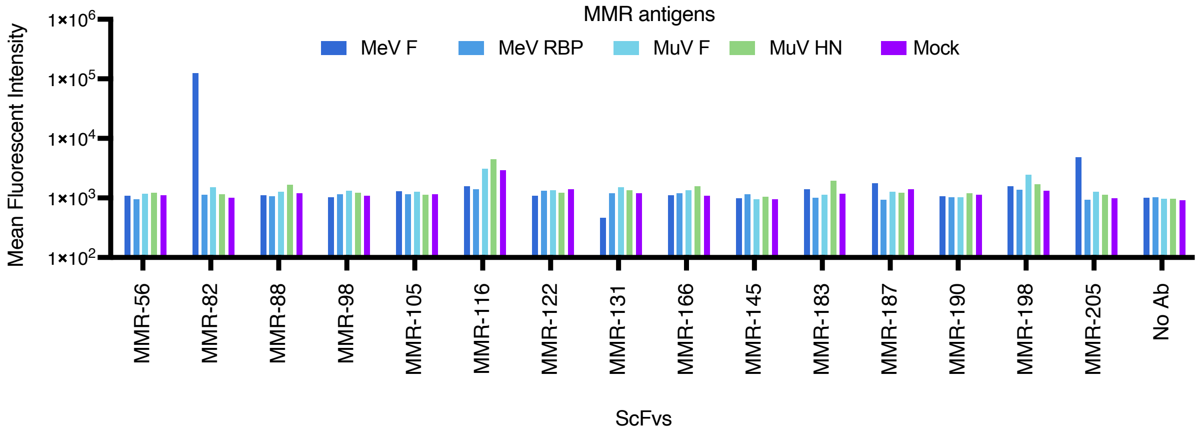

#### Figure S10: V-gene usage in light chains

V-gene distribution in light chains. V-genes were assigned by IgBlast using the personalized database for IGL and the default IMGT database for IGK. As for Figure 6A, sequences that blasted with less than 85% VK or VL identity (light blue) were analyzed separately.

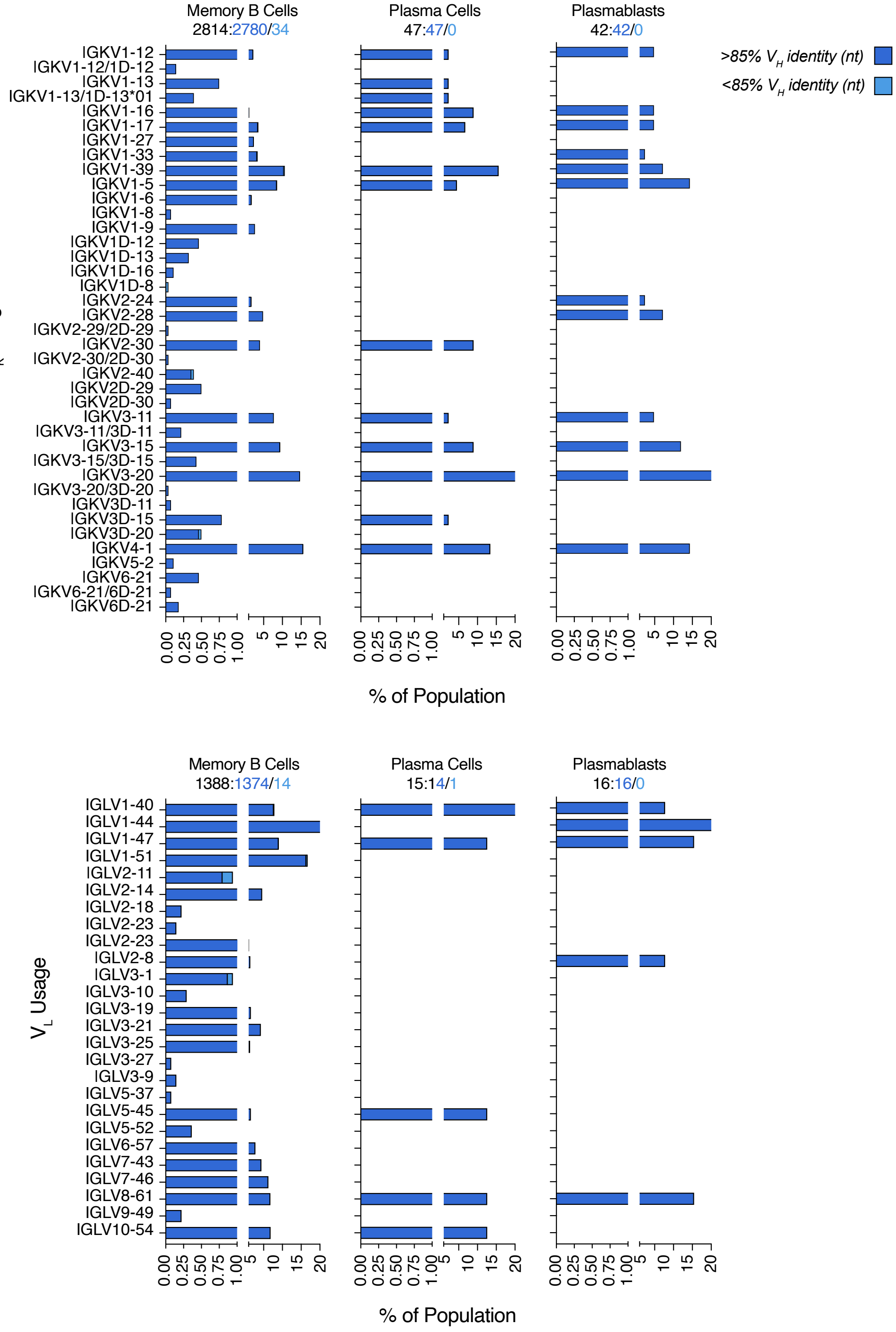

#### Figure S11: J-gene usage

J-gene distribution in light and heavy chains. MG: Ambiguous call between multiple genes.

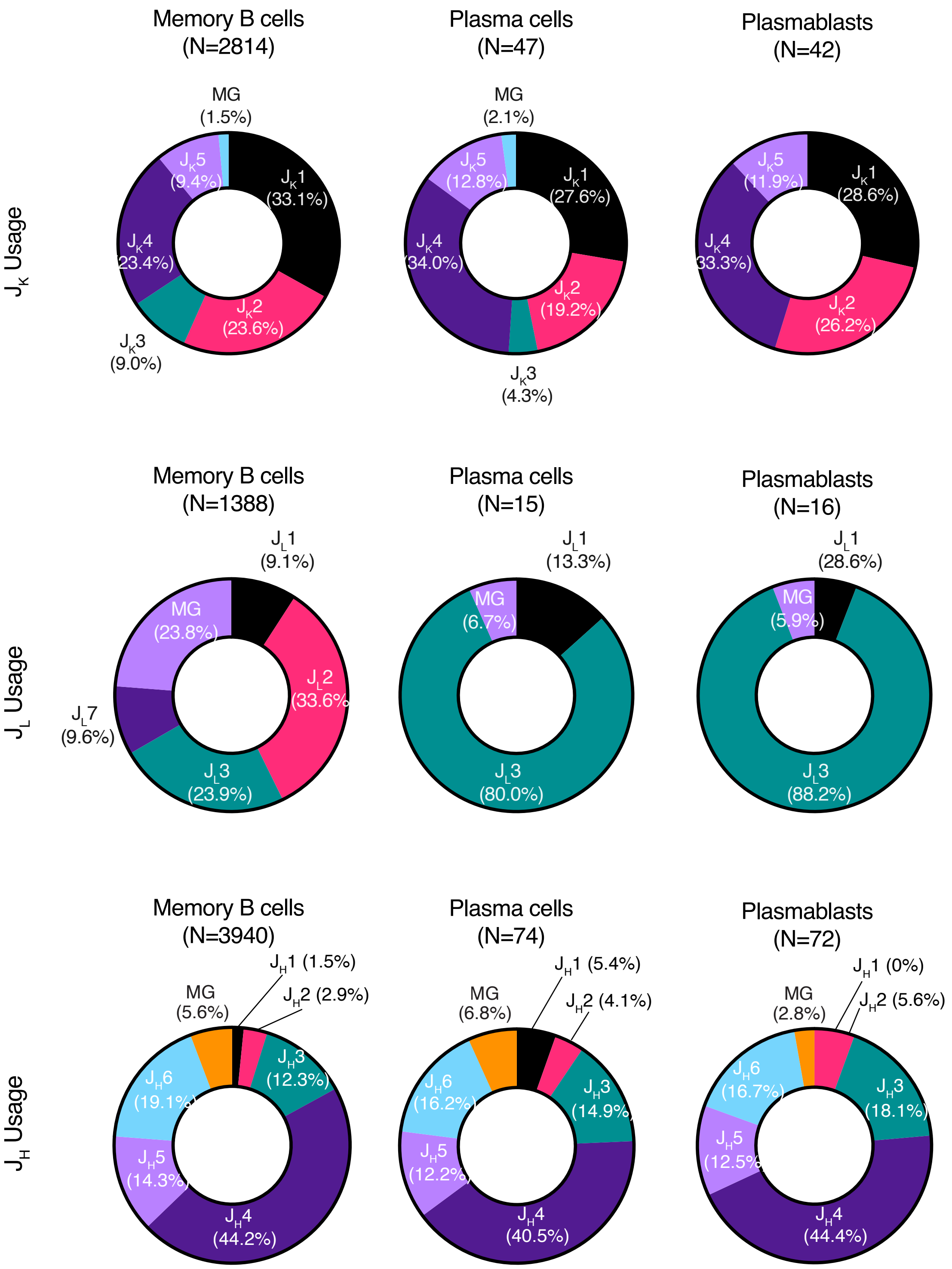

#### Table S1: Whole genome assembly stats

Whole genome assembly statistics from Flye and Margin (as a part of the HapDup pipeline).

| Total Length | Fragments | Fragments N50 | Largest Fragment | Phase Sets | Phase Set N50 | Largest Phase Set | Average Phase Set | Variants | Phased Variants |
|---|---|---|---|---|---|---|---|---|---|
| 2.87 Gb | 2,437 | 28.5 Mb | 109 Mb | 9,791 | 1.0 Mb | 5.4 Mb | 252 Kb | 3,218,774 | 2,766,421 |

#### Table S2: Differentially expressed transcripts in MBC gate

Transcripts whose expression in one cluster was significantly different from the expression in all cells outside of that cluster. Calculated using the FindAllMarkers function in Seurat. Only showing a sample of the table; full table available in Supplemental_Table_S2.xlsx.

| p_val | avg_log2FC | pct.1 | pct.2 | p_val_adj | cluster | gene |
|---|---|---|---|---|---|---|
| 0 | 9.707216369 | 0.88 | 0.002 | 0 | 11 | ENST00000390287 |
| 0 | 8.888899069 | 0.91 | 0.015 | 0 | 7 | ENST00000390310 |
| 0 | 8.796613561 | 0.789 | 0.003 | 0 | 9 | ENST00000468494 |
| 0 | 8.763374157 | 0.688 | 0.002 | 0 | 9 | ENST00000474213 |
| 0 | 8.42241238 | 0.932 | 0.004 | 0 | 8 | ENST00000448155 |
| 0 | 7.735901236 | 0.444 | 0.003 | 0 | 6 | ENST00000390282 |
| 0 | 7.680285344 | 0.96 | 0.017 | 0 | 10 | ENST00000390556 |
| 0 | 6.976460678 | 0.828 | 0.011 | 0 | 10 | ENST00000613640 |
| 0 | 6.84493599 | 0.41 | 0.006 | 0 | 7 | ENST00000390314 |
| 0 | 6.790062411 | 0.934 | 0.038 | 0 | 6 | ENST00000390331 |
| 0 | 6.399653128 | 0.767 | 0.061 | 0 | 4 | ENST00000390298 |
| 2.87E-297 | 6.208927382 | 0.268 | 0.006 | 1.48E-292 | 4 | ENST00000390295 |
| 0 | 5.809345784 | 1 | 0.089 | 0 | 7 | ENST00000390321 |
| 0 | 5.1663246 | 0.938 | 0.046 | 0 | 5 | ENST00000390243 |
| 2.73E-132 | 4.131387271 | 0.983 | 0.197 | 1.41E-127 | 8 | ENST00000498574 |
| 1.07E-47 | 4.068982159 | 0.313 | 0.03 | 5.52E-43 | 11 | ENST00000390285 |
| 5.13E-107 | 3.992857839 | 1 | 0.177 | 2.65E-102 | 11 | ENST00000390309 |
| 3.82E-187 | 3.266694923 | 0.379 | 0.116 | 1.98E-182 | 1 | ENST00000390312 |
| 2.96E-26 | 3.261249593 | 0.317 | 0.151 | 1.53E-21 | 3 | ENST00000460472 |
| 2.01E-28 | 2.653424151 | 0.436 | 0.269 | 1.04E-23 | 3 | ENST00000647309 |
| 0 | 2.64751702 | 0.903 | 0.296 | 0 | 1 | ENST00000390323 |
| 4.67E-12 | 2.6440981 | 0.275 | 0.173 | 2.41E-07 | 3 | ENST00000266022 |
| 9.43E-13 | 2.62822186 | 0.255 | 0.148 | 4.87E-08 | 3 | ENST00000531807 |
| 2.08E-15 | 2.594175249 | 0.303 | 0.181 | 1.07E-10 | 3 | ENST00000622875 |

#### Table S3: Nanopore sequencing statistics

Nanopore sequencing statistics for the single-cell libraries and the germline genome library.

| Library | Sample | Library treatment | Kit | # Flow cells | Flow cell ID | Base calling mode | Total reads (M) | Total yield (Gb) | Read N50 (bp) |
|---|---|---|---|---|---|---|---|---|---|
| Single-cell RNA sequencing | P7 gate: Antibody secreting cells | Whole transcriptome | LSK110 | 2 | PAI05989 | Simplex | 70.4 | 84.3 | 1,336 |
| | | | | | PAI07242 | Simplex | 99.9 | 111.9 | 1,195 |
| | P8 gate: Memory B cells | Whole transcriptome | PCS111 | 2 | PAM80546 | Simplex | 182.1 | 140.7 | 830 |
| | | | | | PAM71314 | Simplex | 187.8 | 144.1 | 825 |
| | | IG-enrichment | LSK110 | 1 | PAI92332 | Simplex | 123.7 | 129.3 | 1,054 |
| Germline genome sequencing | Monocytes | Whole genome | LSK114 | 1 | PAQ76345 | Simplex | 16.9 | 125.4 | 21,173 |
| | | | | | | Duplex | 1.6 | 12.8 | 20,701 |

#### Table S4: Primers for amplification of 10X libraries

Primer sequences used to amplify the 10x single-cell cDNA library prior to sequencing.

| Primer name | Primer Sequence |
|---|---|
| 10x_cDNA_fwd | CTACACGACGCTCTTCCGATCT |
| 10x_cDNA_rev | AAGCAGTGGTATCAACGCAGAG |

#### Table S5: Probe sequences

Probe sequences from the xGen™ Custom Hybridization Panel (Integrated DNA Technologies) targeting BCR and TCR exons. The panel consisted of 1877 5' biotinylated xGen Lockdown Probes, pooled at equimolar concentrations and purified with standard desalting. Only showing a sample of the table; full table available in Supplemental_Table_S5.xlsx.

| Sequence Name | Sequence |
|---|---|
| 783320_45035616_IGKV1OR-2\|exon_2\|ENSG00000156755_1_1 | /5Biosg/GTG CCA GAT GTG ACA TCC AGA TGA CCC AGT CTC CAT CCT CCC TGT CTG CAT CTG TAG GAG GCA GAG TCA CCA TCA CTT GCC GGG CGA GTC AGG GCA TTA GCA ATA ATT TAA ATT GGT ATC |
| 783320_45035616_IGKV1OR-2\|exon_2\|ENSG00000156755_1_2 | /5Biosg/CAG GGC ATT AGC AAT AAT TTA AAT TGG TAT CAG CAG AAA CCA AGG AAA ACT CCT AAG CTC CTG ATC TAT GCT GCA TCC AGT CTG CAA AGT GGG ATT CCC TCT CGG TTC AGT GAC AGT GGA |
| 783320_45035616_IGKV1OR-2\|exon_2\|ENSG00000156755_1_3 | /5Biosg/TGG GAT TCC CTC TCG GTT CAG TGA CAG TGG ATC TGG GAC AGA TTA CAC TCT CAC CAT CAG CAG CCT GCA GCC TGA AGA TTT TGC AAC TTA TTA CTG TCA ACA GAG TGA CAG TAA CCC TCC |
| 783320_45035617_TRBV21OR9-2\|exon_1\|ENSG00000183938_1_1 | /5Biosg/CCT CCA TGG ACA CCA AGG TCA CCC AGA GAC CTA GAT TTC TGG TCA AAG CAA ATG AAC AGA AAG CAA AGA TGG ACT GTG TTC CTA TAA AAA GAC ATA GTT ATG TTT ACT GGT ATC ATA AGA |
| 783320_45035617_TRBV21OR9-2\|exon_1\|ENSG00000183938_1_2 | /5Biosg/AAG ACA TAG TTA TGT TTA CTG GTA TCA TAA GAC GCT GGA AGA AGA GCT CAA GTT TTT TAT TTA CTT TCA GAA TGA AGA AAT TAT TCA GAA AGC AGA AAT AAT CAA TGA GCG ATT TTC AGC |
| 783320_45035617_TRBV21OR9-2\|exon_1\|ENSG00000183938_1_3 | /5Biosg/AAA GCA GAA ATA ATC AAT GAG CGA TTT TCA GCC CAA TGC CCC CAA AAC TCA CCC TGT ACC TTG GAG ATC CAG TCC ACG GAG TCA GGA GAC ACA GCA CGG TAT TTC TGT GCC AAC AGC AAA |
| 783320_45035618_IGHV1OR15-9\|exon_2\|ENSG00000188403_1_1 | /5Biosg/GTG CCC AGT CCC AGG TAC AGC TGA TGC AGT CTG GGG CTG AGG TGA AGA AGC CTG GGG CCT CAG TGA GGA TCT CCT GCA AGG CTT CTG GAT ACA CCT TCA CCA GCT ACT GTA TGC ACT GGG |
| 783320_45035618_IGHV1OR15-9\|exon_2\|ENSG00000188403_1_2 | /5Biosg/CTT CAC CAG CTA CTG TAT GCA CTG GGT GTG CCA GGC CCA TGC ACA AGG GCT TGA GTG GAT GGG ATT GGT GTG CCC TAG TGA TGG CAG CAC AAG CTA TGC ACA GAA GTT CCA GGG CAG AGT |
| 783320_45035618_IGHV1OR15-9\|exon_2\|ENSG00000188403_1_3 | /5Biosg/TAT GCA CAG AAG TTC CAG GGC AGA GTC ACC ATA ACC AGG GAC ACA TCC ATG GGC ACA GCC TAC ATG GAG CTA AGC AGC CTG AGA TCT GAG GAC ACG GCC ATG TAT TAC TGT GTG AGA GA |
| 783320_45035619_IGKV7-3\|exon_1\|ENSG00000197794_1_1 | /5Biosg/GGG ACA TTG TGC TGA CCC AGT CTC CAG CCT CCT TGG CCG TGT CTC CAG GAC AGA GGG CCA CCA TCA CCT GCA GAG CCA GTG AGA GTG TCA GTT TCT TGG GAA TAA ACT TAA TTC ACT GGT |
| 783320_45035619_IGKV7-3\|exon_1\|ENSG00000197794_1_2 | /5Biosg/AGT TTC TTG GGA ATA AAC TTA ATT CAC TGG TAT CAG CAG AAA CCA GGA CAA CCT CCT AAA CTC CTG ATT TAC CAA GCA TCC AAT AAA GAC ACT GGG GTC CCA GCC AGG TTC AGC GGC AGT |
| 783320_45035619_IGKV7-3\|exon_1\|ENSG00000197794_1_3 | /5Biosg/CAC TGG GGT CCC AGC CAG GTT CAG CGG CAG TGG GTC TGG GAC CGA TTT CAC CCT CAC AAT TAA TCC TGT GGA AGC TAA TGA TAC TGC AAA TTA TTA CTG TCT GCA GAG TAA GAA TTT TCC |
| 783320_45035620_IGKV1OR2-3\|exon_1\|ENSG00000204670_1_1 | /5Biosg/TGA CAT CCA GAT GAC CCA GCC TCC ATC CTC CCT GTC TGC ATC TGT AGG AGA CAG AGT CAC CGT CTC TTG CCA GGC TAG TCA AAG CAT TTA CAA CTA TTT AAA TTG GTA TCA GCA GAA ACC |
| 783320_45035620_IGKV1OR2-3\|exon_1\|ENSG00000204670_1_2 | /5Biosg/CAT TTA CAA CTA TTT AAA TTG GTA TCA GCA GAA ACC AGG GAA AGC ACC TAA GTT CCT GAC CTA TAG GGC ATC CAG TTT GCA GAG GGG GAT GCC ATC TCA GTT CAG TGG CAG CGG ATA TGG |
